# Epigenetic Reprogramming Alters Human Intestinal Stem Cell Fate in Pouchitis

**DOI:** 10.1101/2025.03.19.644201

**Authors:** Chaoting Zhou, Shan Liu, Jing Yu Carolina Cen Feng, Katherine S. Ventre, Tianyi Li, Jordan E. Axelrad, Kyung Ku Jang, Ken Cadwell

**Affiliations:** Cell and Molecular Biology Graduate Program, University of Pennsylvania Perelman School of Medicine, Philadelphia, PA 19104, USA; Division of Gastroenterology and Hepatology, Department of Medicine, University of Pennsylvania Perelman School of Medicine, Philadelphia, PA 19104, USA; Center for Cellular Immunotherapies, Perelman School of Medicine at the University of Pennsylvania, Philadelphia, PA 19104, USA; Department of Biomedical Engineering, NYU Tandon School of Engineering, Brooklyn, NY, 11201 USA; Division of Gastroenterology and Hepatology, Department of Medicine, New York University Grossman School of Medicine, New York, NY 10016, USA; Ronald O. Perelman Department of Dermatology, New York University Grossman School of Medicine, New York, NY 10016, USA; Master of Biotechnology Program, University of Pennsylvania School of Engineering and Applied Science, Philadelphia, PA 19104, USA; Department of Microbiology, New York University Grossman School of Medicine, New York, NY 10016, USA; Department of Anatomy, Graduate School of Medical Science, Brain Korea 21 Project, Yonsei University College of Medicine, Seoul 03722, South Korea; Department of Pathobiology, University of Pennsylvania Perelman School of Veterinary Medicine

## Abstract

Intestinal stem cells (ISCs) mediate the continuous renewal of the epithelium during homeostasis and recovery from injury. To investigate the molecular impact of inflammation on human ISCs and the role of janus kinase/signal transducer and activator of transcription (JAK/STAT) signaling, we established an organoid model derived from individuals with pouchitis, a form of inflammatory bowel disease that develops in an organ generated from ileal pouch-anal anastomosis (IPAA) surgery. Compared with non- inflamed specimens, pouchitis organoids exhibited increased apoptosis and secretory lineage differentiation that were stable after multiple passages, mirroring findings in primary pouch tissue. Chromatin accessibility and histone modification profiling of ISCs revealed inflammation-driven epigenetic remodeling, particularly involving activator protein 1 (AP-1) transcription factors such as c- Jun and excessive STAT1 activity. Loss of c-Jun disrupted ISC viability and enhanced secretory cell differentiation in non-inflamed organoids, whereas therapies targeting JAK/STAT reversed these ISC defects and chromatin changes in pouchitis organoids. Together, these results associated inflammation with epigenetic changes in human ISCs, suggesting that immune-mediated injury has lasting effects on the pouch epithelium that can potentially be reversed through JAK/STAT therapies.

**One Sentence Summary:** Persistent epigenetic remodeling in the epithelium during pouchitis is enforced by excessive JAK/STAT activity.

## INTRODUCTION

Intestinal stem cells (ISCs) are responsible for the continuous renewal of the epithelium, ensuring proper barrier and digestive function over the entire lifespan (*1*, *2*). Located at or near the base of the crypts, ISCs differentiate into absorptive enterocytes and secretory cell types such as goblet, enteroendocrine, and Paneth cells that produce mucus, hormones, and antimicrobials, respectively. The remarkable self- renewing capacity of the epithelium is illustrated by the ability of ISCs to form epithelial organoids by differentiating and self-organizing into three-dimensional structures when cultured with growth factors and an extracellular matrix. Intestinal organoids replicate certain aspects of the tissue from which they are generated and have facilitated the discovery of extracellular and intracellular cues that dictate epithelial self-renewal and differentiation, such as the critical role of WNT-β-catenin and NOTCH signaling that work together with transcription factors (TFs) and chromatin modifications to preserve ISC plasticity (*3–6*). Also, findings from human organoids and mouse models have supported each other by showing how ISC survival and differentiation can be further impacted by immune mediators, such as the lymphocyte cytokines interferon-γ (IFN-γ) and interleukin-17 (IL-17) (*7–12*), underscoring the responsiveness of the epithelium to the immune environment.

Chronic immune-mediated damage to the intestinal epithelial barrier is a hallmark of several major disorders including inflammatory bowel disease (IBD), graft-versus-host disease (GVHD), and Celiac disease. Following wounding, ISCs rapidly proliferate and differentiate into new cells that replace the damaged epithelium and return to their original state. However, recovery may be more complicated in the setting of chronic inflammation. Intestinal biopsies obtained from patients with IBD generally display reduced capacity to form viable organoids, suggesting damage to ISCs (*13*, *14*). In mice, tissue stem cells retain a memory of inflammation, influencing their subsequent fate and function. For example, signal transducer and activator of transcription 3 (STAT3) cooperates with the activator protein 1 (AP-1) TFs FOS and JUN to mediate chromatin accessibility in regions associated with lineage plasticity in skin epidermal stem cells to enhance wound repair during a recall response (*15*). Given that the janus kinase/signal transducer and activator of transcription (JAK/STAT) pathway mediate cytokine receptor signaling, this observation provides a direct link between the external inflammatory environment and the chromatin landscape of the stem cell compartment. In contrast to the skin, alterations to the epigenome of ISCs in a mouse model of GVHD lead to adverse reductions in differentiation and the ability to form organoids (*16*). The impact of inflammation on human ISCs and the underlying mechanism are less clear.

IBD comprises two major subtypes: ulcerative colitis (UC), which is limited to the colon and rectum, and Crohn’s disease (CD), which can affect any part of the gastrointestinal tract. Surgical removal of affected regions, including proctocolectomy, is often required in patients with medically refractory disease. For these patients, ileal pouch-anal anastomosis (IPAA) is a continence-preserving staged surgical procedure that enhances quality of life by eliminating the need for a permanent ostomy. After removal of the colon and rectum, a loop of ileum is sewn into a J-shaped pelvic reservoir (J-pouch) that is subsequently attached to the anus. Although the J-pouch is created from non-inflamed ileal tissue, up to 50% of patients with IBD experience a de novo inflammatory disease known as pouchitis in this surgically created organ, with nearly 70% of cases occurring within the first year following IPAA surgery (*17–19*). The mechanism of pouchitis is unknown but is rare in patients without IBD who undergo an IPAA. We and others have shown that the immune landscape of pouchitis resembles IBD and includes infiltration by inflammatory T cells and myeloid cells and adoption of a pro-inflammatory transcriptional profile (*20–22*). As with other forms of IBD, JAK inhibitors are emerging as therapeutic agents for pouchitis refractory to conventional therapies (*23*). The J-pouch epithelium, particularly enterocytes, undergo partial colonic metaplasia in patients with UC, suggesting a link between epithelial differentiation and pouchitis (*24*). Given the rapid post-surgical onset of disease in individuals with IBD who undergo IPAA, pouchitis provides a unique opportunity to investigate the impact of inflammation and therapies targeting JAK/STAT on the ISC compartment.

In this study, we established a pouch-derived organoid model to investigate the pathophysiology of pouchitis. Previous studies using patient-derived intestinal organoids from individuals with IBD have shown that disease-associated epithelial characteristics can persist ex vivo, such as increased antimicrobial gene expression and reduced differentiation of goblet cells, although the precise defects differ between cohorts (*25–28*). Also, organoids generated from children with IBD display distinct and stable DNA methylation patterns that differ between disease subsets and affected region of the gut (*29*), highlighting the need for a pouchitis-specific examination. To address this gap, we examined ISC properties and associated chromatin and transcriptional states in organoids-derived from the J-pouch of patients with and without pouchitis.

## RESULTS

### ISCs from the inflamed J-pouch display reduced viability via caspase-mediated apoptosis

J-pouch organoids have not been previously described. The pouch body (distal reservoir region) is exposed to prolonged fecal stasis, likely increasing exposure to microbes and cytokines, whereas the pouch inlet (distal to the neo-terminal ileum) is anatomically and functionally closer to the native ileum. These differences may contribute to distinct epithelial properties. Therefore, we analyzed both pouch body- and inlet-derived organoids from patients with a history of UC with or without inflammation (Fig. 1A).

**Fig. 1.**
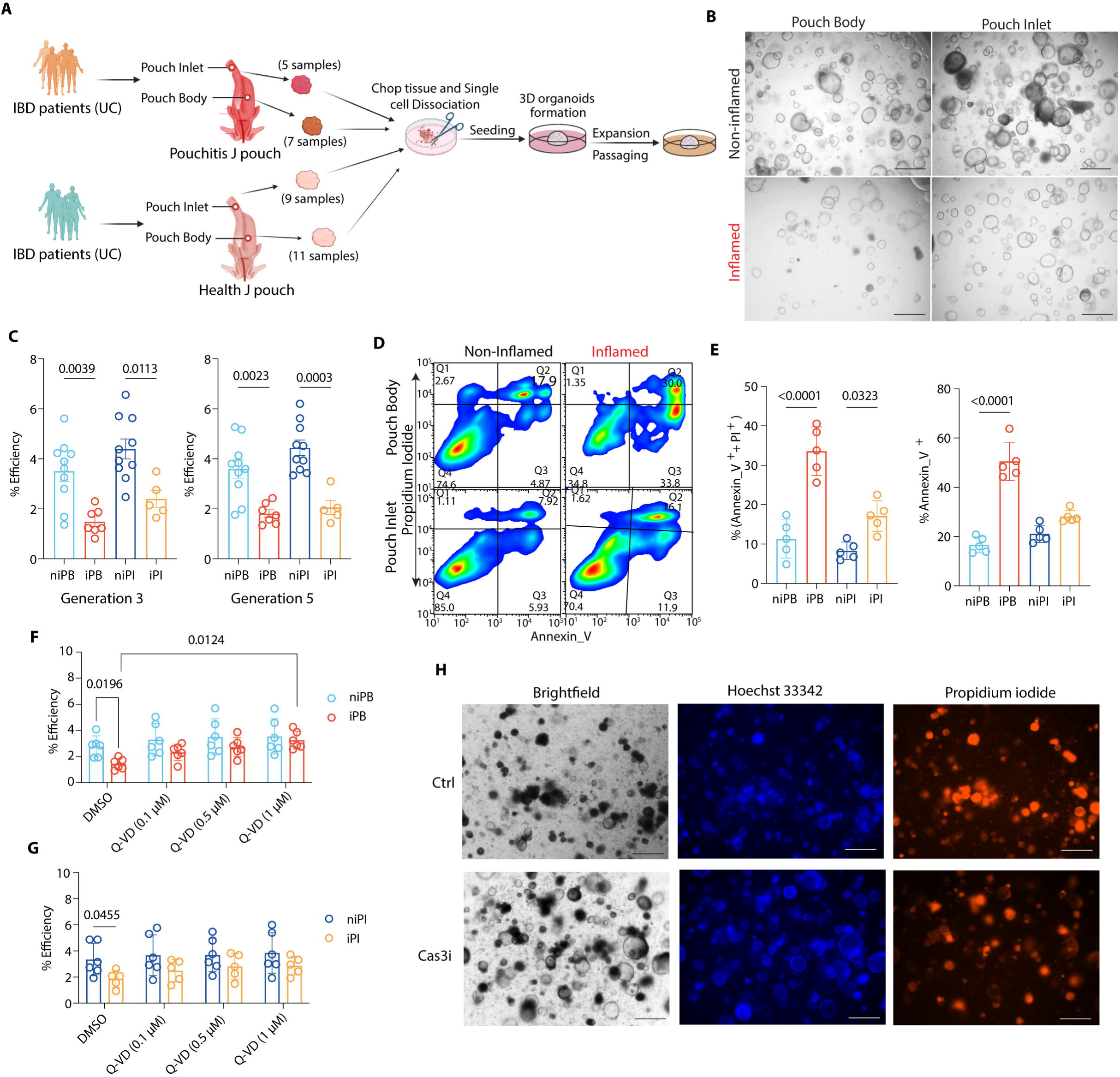
ISCs from the inflamed J-pouch display reduced viability. **(A)** Experimental design for J-pouch tissue collection from patients with ulcerative colitis (UC), with and without pouchitis. **(B and C)** Representative images (B) and quantification of generation efficiency (C) of organoids derived from unaffected regions of the non-inflamed pouch body (niPB) and inlet (niPI) or inflamed (pouchitis) pouch body (iPB) and inlet (iPI). Scale bars, 400 μm. **(D and E)** Representative flow cytometry plots (D) and quantification (E) of ISCs gated on CD24^+^CD166^+^CD44^+^PTK7^high^ cells in organoids from niPB, niPI, iPB, and iPI stained for propidium iodide (PI) and annexin V. ISC gating strategy in Figure S1A. **(F and G)** Quantification of generation efficiency of organoids derived from niPB, iPB, niPI, and iPI treated with vehicle (DMSO) or indicated concentrations of Q-VD-OPh. **(H)** Images of iPB organoids treated with DMSO, 10 μM caspase-3 inhibitor (Cas3i, Z-DEVD-FMK), visualized by brightfield microscopy, Hoechst 33342, and PI staining. Scale bars, 400 μm. Dots represent individual samples; bars show mean ± SD. Statistical significance was determined by one-way ANOVA with Tukey’s multiple-comparisons test (C and E) or two-way ANOVA with Holm–Šidák correction for multiple comparisons (F and G). Exact *P* values are shown.

Organoids were successfully generated and expanded from these pouch specimens using previously described conditions for organoids of colonic origin (detailed in Methods) (*30*) and passaged three generations before initiating experiments to ensure that observations represent stable properties. The generation efficiency of organoids from inflamed tissue was less efficient when compared to organoids derived from non-inflamed tissue, even after multiple passages and freeze-thaw cycles (fig. S1A). To enable rigorous quantification of generation efficiency and assessment of differentiation, single cell suspensions were generated from passaged organoids on day 7 post-seeding, and each well of a 96 well plate was seeded with 3000 cells in triplicate to standardize the initial cell number and maintained in differentiation media containing insulin-like growth factor 1 (IGF-1) and fibroblast growth factor 2 (FGF-2) (*31*) (Fig. S1B). Culturing in differentiation media for 7 days led to upregulation of secretory lineage and Paneth cell marker *secretory phospholipase 2* (*sPLA2*), secretory cell lineage TF *ATOH1*, enteroendocrine cell marker *chromogranin A* (*CHGA*), goblet cell marker *mucin 2* (*MUC2*), and enterocyte markers *fatty acid binding protein 2* (*FABP2*) and *apolipoprotein A1* (*APOA1*) (fig. S1C). These initial analyses indicated that organoids can be generated from pouch inlet and body biopsies, serially passaged, and cultured in previously described expansion and differentiation media.

Compared with organoids from non-inflamed pouch inlet (niPI) and pouch body (niPB), we observed a marked reduction in the organoid formation efficiency for those that were derived from inflamed pouch inlet (iPI) and pouch body (iPB) (Fig. 1, B and C). Ki-67 staining, a marker of cell division, did not reveal a reduction in proliferating epithelial cells in iPB compared with niPB (fig. S1, D and E). We therefore examined whether increased cell death contributed to the reduced establishment of inflamed pouch organoids. In the colon, human ISCs were identified as CD24^+^CD166^+^CD44^+^PTK7^high^ cells that are sufficient to form organoids (*32*, *33*). These cells were detected in J-pouch tissue by flow cytometry (fig. S1, F and G). Fluorescence-activated cell sorting (FACS)-sorted CD24^+^CD166^+^CD44^+^PTK7^high^ (Population 3; P3) cells formed organoids at highest efficiency, while CD24^+^CD166^+^CD44^+^PTK7^low^ (P2) formed organoids at lower frequencies and CD24^-^CD44^+^ (P1) cells failed to form organoids entirely (fig. S1, H and I). iPB and iPI organoids generated from sorted P3 cells displayed reduced formation efficiency and were less viable by MTT staining compared with niPB and niPI organoids (fig. S1, H and I). Flow cytometry analysis showed increased proportions of ISCs that were Annexin V^+^ propidium iodide (PI^+^) or Annexin V^+^ PI^-^ from iPB and iPI organoids compared with their non-inflamed counterparts, indicating higher rates of apoptosis in the inflamed group (Fig. 1, D and E). Blocking apoptosis with the pan-caspase inhibitor Q-VD-OPh improved organoid formation efficiency in inflamed samples to levels similar to non-inflamed samples (Fig. 1, F and G). Western blotting (WB) revealed an increase in cleaved PARP and caspase-3, -7, and -9 to various degrees, but not caspase-8, in iPB and iPI organoids cultured for 4 hours in differentiation medium (fig. S2, A and B), consistent with activation of the intrinsic pathway of apoptosis. Chemical inhibitors of caspase-3 and caspase-9 improved iPB and iPI organoid formation efficiency whereas caspase-8 inhibition and ER stress inhibitor taurosodeoxycholic acid (TUDCA) did not rescue the impaired organoid formation (Fig. 1H, and fig. S2, C and E). These results indicated that organoids from pouchitis specimens display persistent deficits in generation efficiency due to ISC apoptosis.

### Pouchitis organoids display enhanced differentiation of secretory epithelial cells

To determine whether the inflammatory state of the pouch affects ISC differentiation, we analyzed cell lineage markers by immunofluorescence microscopy on day 10 of organoid culture in differentiation media. We found that the number of CHGA^+^ and MUC2^+^ cells were increased and APOA1^+^ cells were decreased in iPB compared to niPB organoids (Fig. 2A to D). iPI organoids exhibited similar trends when compared with niPI organoids, but the differences did not reach statistical significance (Fig. 2B, and fig. S3A). The number of cells positive for the Paneth cell marker lysozyme (LYZ) were similar across all groups (Fig. 2E). Transcriptional analysis by RT-qPCR largely supported these findings. Enteroendocrine cell transcripts *CHGA*, *NEUROD1*, and *NEUROD3* displayed higher expression in iPB organoids compared with others, whereas *APOA1* and *HES1* were decreased. When we examined primary J-pouch tissue sections from the same patients from whom biopsies were used to generate organoids, iPB tissue contained more CHGA cells and MUC2 cells and fewer APOA1 cells than niPB tissue (Fig. 2, G and H). Similar trends were observed in pouch inlet, though differences were not statistically significant (Fig. 2G, and fig. S3B). LYZ cells, which were located in the crypts, were lower in number in iPB compared with niPB tissue (Fig. 2, G and H). Thus, the staining patterns for CHGA, MUC2, and APOA1 epithelial markers in vivo resembled our findings in organoids. Overall, these findings indicated that ISCs from the inflamed J-pouch are skewed toward differentiation into enteroendocrine and goblet cells both in vivo and in vitro.

**Fig. 2.**
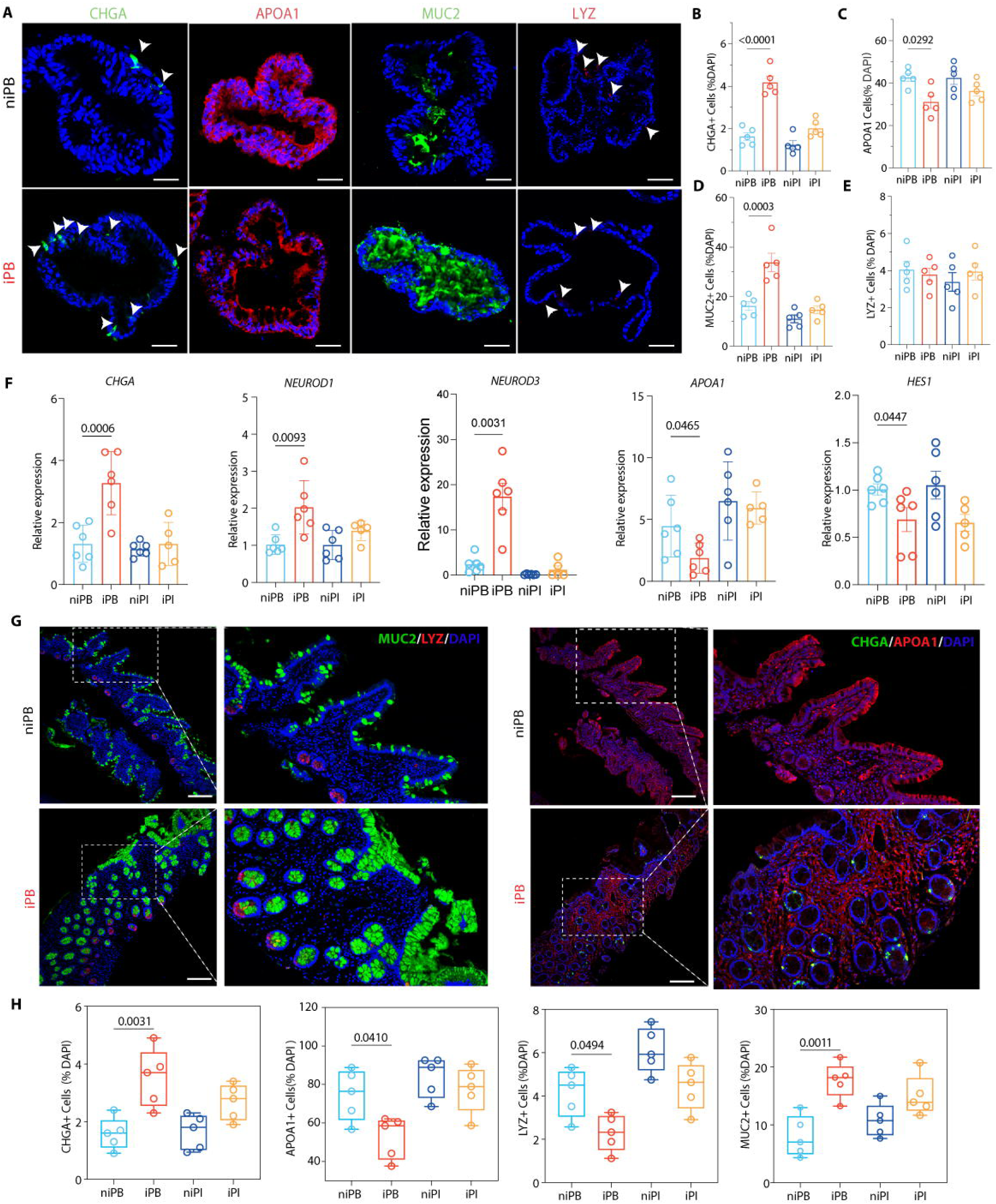
Pouchitis organoids display enhanced differentiation of secretory epithelial cells. **(A)** Immunofluorescence staining of organoids generated from inflamed and non-inflamed pouch body and inlet for MUC2, LYZ, CHGA, APOA1. Scale bars, 200 µm. **(B - E)** Quantification of the proportion of CHGA^+^ (B), APOA1^+^ (C), MUC2^+^ (D), and LYZ^+^ (E) cells normalized to the total number of cells based on DAPI in organoids from (A). **(F)** Quantitative real-time PCR (RT-qPCR) analysis of *CHGA*, *NEUROD3, NEUROD1*, *APOA1* and *HES1* expression in organoids derived from non-inflamed and inflamed tissues of the J-pouch normalized to *GAPDH*. **(G)** Immunofluorescence microscopy of unaffected J-pouch tissue sections from individuals with or without pouchitis stained for CHGA, APOA1, LYZ, and MUC2. Scale bars, 50 µm. **(H)** Quantification of MUC2^+^, LYZ^+^, CHGA^+^, and APOA1^+^ cells from (G). Dots represent individual samples; bars show mean ± SD; boxes show median and interquartile range, with whiskers indicating min–max. Statistical significance was determined by one-way ANOVA with Tukey’s post hoc test. Exact *P* values are shown.

### Inflammation alters the chromatin landscape of J-pouch ISCs

Because differences due to the inflammatory status of the original tissue persisted through multiple passages of organoid culture, we hypothesized that inflammation induces stable epigenetic alterations within ISCs. To investigate this possibility, we performed assay for transposase-accessible chromatin using sequencing (ATAC-seq) on FACS-isolated ISCs derived from inflamed and non-inflamed pouch organoids. Principal Component Analysis (PCA) revealed that ISCs from iPB displayed the most distinct pattern (fig. S4, A and B). We observed extensive remodeling of chromatin accessibility in iPB ISCs, characterized by widespread gains and losses of accessibility across regulatory regions compared with other groups (Fig. 3, A and B). In contrast, chromatin accessibility changes were substantially less pronounced in iPI ISCs, indicating that inflammation-associated epigenetic remodeling is more prominent in the pouch body than in the pouch inlet (fig. S4, C to E).

**Fig. 3.**
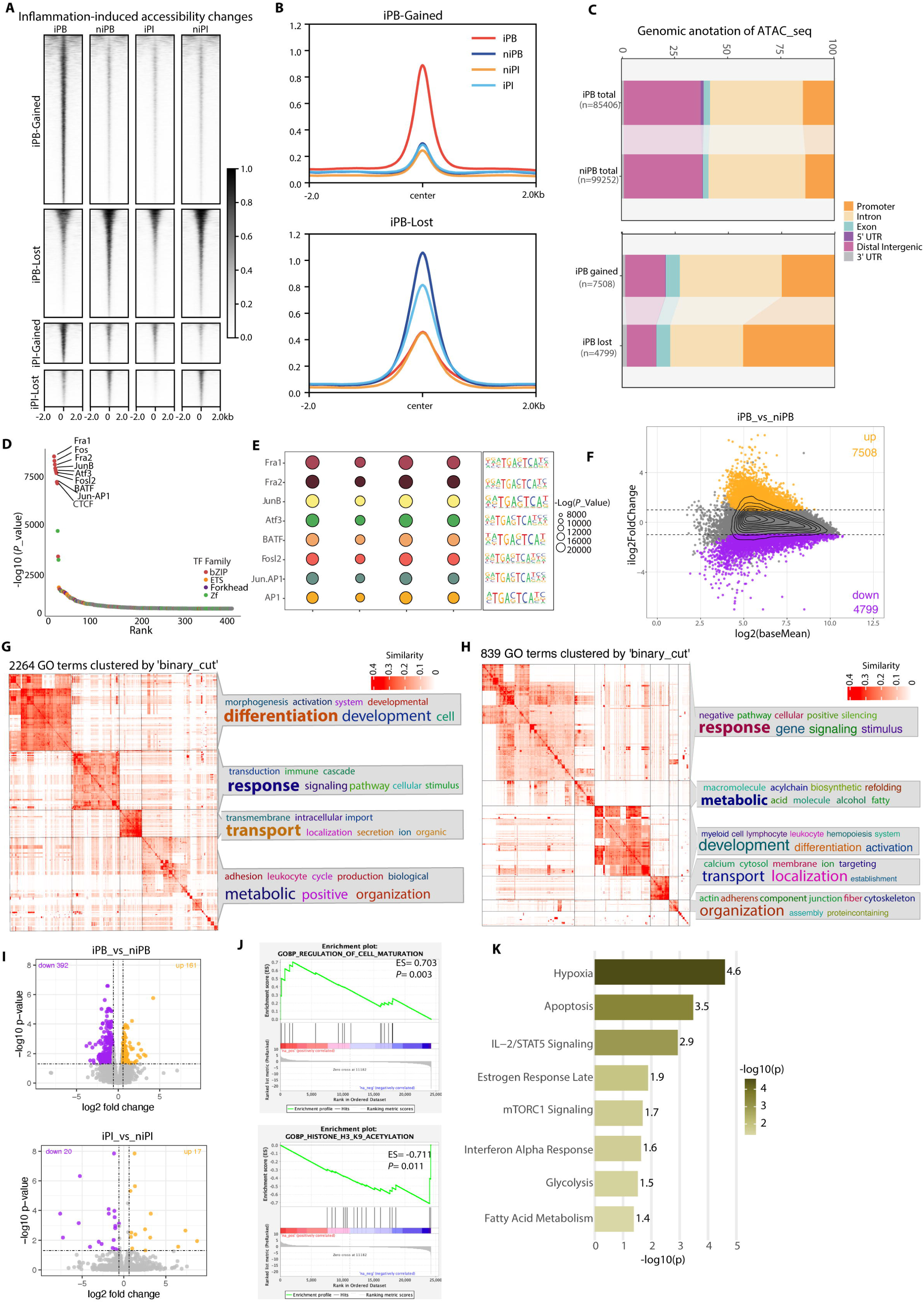
Inflammation alters the chromatin landscape of J-pouch ISCs. **(A and B)** Heatmaps (A) and average signals plots (B) across regions gaining or losing accessibility centered on peak summits (±2 kb) identified by ATAC-seq of ISCs sorted from iPB, niPB, iPI, and niPI organoids (n = 3-4 each). **(C)** Genomic annotation of accessible regions in iPB and niPB and regions gained or lost in iPB versus niPB. **(D and E)** Transcription factor motifs enriched in accessible regions (D) and comparison across groups (E). **(F)** MA plot of regions with gained or lost chromatin accessibility in iPB versus niPB determined by DESeq2 (*P_adj_* < 0.05, |FC| ≥ 1.5). **(G and H)** Semantic clustering of Gene Ontology (GO) biological process terms associated with chromatin regions with gained (G) or lost accessibility (H) in iPB versus niPB ISCs. GO terms were identified by GREAT analysis and clustered using the binary-cut algorithm. **(I)** Volcano plots showing differentially expressed genes according to RNA-seq analysis comparing iPB versus niPB or iPI versus niPI determined by DESeq2 (*P_adj_* < 0.05, |FC| ≥ 1.5). **(J)** Gene set enrichment analysis (GSEA) of differentially expressed genes identified by RNA-seq comparing iPB and niPB ISCs. **(K)** Top enriched MSigDB_Hallmark_2020 pathway based on the -log10(*P*-value).

Differentially accessible regions were predominantly located within promoter and intronic regions (Fig. 3C, fig. S4D). To identify TF networks associated with these chromatin alterations, we performed motif enrichment analysis, which detects transcription factor recognition sequences within accessible chromatin regions but does not directly infer transcription factor binding or activity. AP-1 family motifs belonging to the basic leucine zipper (bZIP) TF family were highly represented across accessible chromatin regions in all groups (Fig. 3D), indicating that AP1 associated cis-regulatory elements are a prominent feature of the pouch ISC chromatin landscape. Comparison across conditions revealed that AP1/JUN associated motif enrichment was lowest in iPB ISCs (Fig. 3E). These findings suggested that inflammation is associated with suppression of AP-1-dependent regulatory programs.

Differential accessibility analysis identified 7,508 regions with increased accessibility and 4,799 regions with decreased accessibility in iPB relative to niPB ISCs (Fig. 3F). In contrast, only 58 gained and 68 lost regions were detected in iPI relative to niPI ISCs (Fig. S5A). Gene ontology analysis of regions exhibiting altered accessibility in iPB ISCs revealed enrichment of biological processes associated with epithelial differentiation, developmental programs, cellular responses to environmental stimuli, transport, and metabolic regulation (Fig. 3, G and H). Regions gaining accessibility in iPB were enriched for stress-response and inflammatory signaling pathways, whereas regions losing accessibility were preferentially associated with epithelial development, morphogenesis, and differentiation programs, suggesting a shift from epithelial homeostatic programs toward inflammatory transcriptional states. In comparison, GO analysis of the smaller set of altered regions in iPI ISCs showed more restricted enrichment, with the most prominent associations related to metabolic processes, particularly biosynthetic and catabolic pathways, along with actin organization and cell-communication programs (fig. S5, E and F). Together, these findings suggested that inflammation induces broad remodeling of epithelial regulatory programs in the pouch body compartment.

We next performed a direct comparison between pouch body and inlet regions. In contrast to the minimal differences between niPB and niPI, we identified 4,311 regions with increased accessibility and 1,602 regions with decreased accessibility in iPB compared with iPI (fig. S5B). Only a small fraction of inflammation-associated regions was shared between the two anatomical compartments (fig. S5, C and D). The pouch body-specific changes were associated with pathways involved in epithelial differentiation, development, apoptosis, and responses to external stimuli (fig. S5, E and F).

RNA sequencing (RNA-seq) of sorted ISCs revealed 392 genes down-regulated and 161 genes up- regulated in iPB relative to niPB, whereas only 37 genes showed significant differential expression between niPI and iPI, mirroring the more limited chromatin level changes observed by ATAC-seq (Fig. 3I). We observed a significant positive correlation between chromatin accessibility and gene expression changes (Spearman’s ρ = 0.586, *P* = 6.62 × 10^-6^, fig. S6A). The correlation at the genome-wide level was modest, suggesting that chromatin alterations act in conjunction with other regulatory mechanisms or may be more important once ISCs differentiate. Concordant alterations in accessibility and expression were observed for genes involved in epithelial differentiation, inflammatory signaling, and metabolic regulation (fig. S6, B and C). Gene set enrichment analysis (GSEA) of transcriptional changes in ISCs associated with inflammation indicated alterations to pathways involved in histone modification, fatty acid metabolism, and toll-like receptor signaling (Fig. 3J, fig. S6D). Similar analysis of inflammation-associated chromatin regions in iPI ISCs identified stress-response, metabolic, homeostatic, and cell-communication pathways, although these changes were much more limited than those observed in iPB ISCs (fig. S4, F and G). Pathway analysis further revealed enrichment of apoptosis, mTORC1 signaling, interferon responses, oxidative phosphorylation, and related inflammatory/stress programs in iPB compared with niPB ISCs (Fig. 3K). These findings demonstrated that pouch body ISCs originating from an inflamed organ display extensive chromatin remodeling accompanied by transcriptional alterations.

### Altered enhancer accessibility in ISCs are associated with H3K27ac modification

We assessed histone H3 lysine 4 monomethylation (H3K4me1) and acetylation of histone 3 lysine 27 (H3K27ac) at cis-regulatory elements on ISCs sorted from organoids using cleavage under targets and release using nuclease (CUT&RUN). H3K4me1 serves as a marker of enhancer priming, indicating that a region is poised for activation, whereas H3K27ac marks active enhancers, reflecting ongoing transcriptional activity. Enhancers that overlap with H3K27ac peaks were classified as active enhancers, and a total enhancer catalog based on H3K4me1 was generated by excluding regions marked with H3K27ac. We first examined histone modification patterns across regions that gained or lost accessibility in iPB ISCs. Regions exhibiting increased accessibility in iPB showed corresponding increases in H3K27ac signal, whereas regions losing accessibility displayed reduced H3K27ac enrichment (Fig. 4A). In contrast, H3K4me1 signals were largely similar between conditions across both gained and lost accessibility regions (Fig. 4A), suggesting that inflammation preferentially affects enhancer activation rather than enhancer priming. Consistent with these observations, differential CUT&RUN analysis identified 1,899 regions with increased H3K27ac and 3,201 regions with decreased H3K27ac in iPB relative to niPB ISCs, whereas only three gained and two lost H3K4me1 regions were detected (Fig. 4B). PCA further demonstrated clear separation of iPB and niPB samples for H3K27ac and not H3K4me1 profiles (fig. S7, A and B). To systematically define enhancer states, we integrated H3K27ac, H3K4me1, and ATAC-seq datasets. Among 79,056 accessible enhancers identified by overlapping ATAC-seq and H3K4me1 peaks, 34,438 were classified as active enhancers based on the presence of H3K27ac (Fig. 4C). Of these active enhancers, 5,100 exhibited differential H3K27ac marking between iPB and niPB ISCs, indicating widespread remodeling of enhancer activity.

**Fig. 4.**
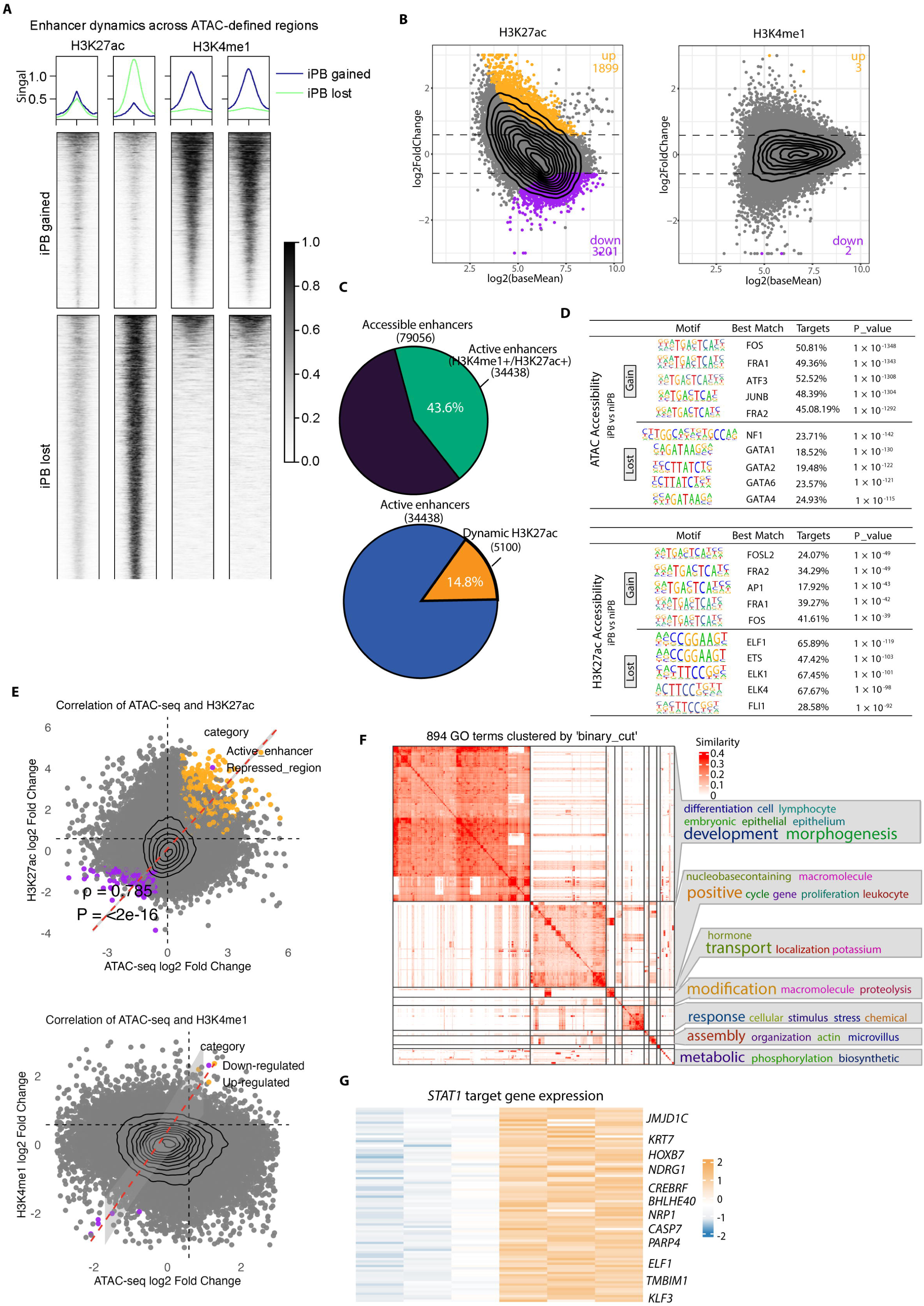
Altered enhancer accessibility in inflamed pouch body ISCs is associated with H3K27ac remodeling. **(A)** Heatmaps of H3K27ac and H3K4me1 distribution in all accessible chromatin regions in ISCs, centered on peak summits (±2 kb). **(B)** MA plots showing accessible enhancers differentially marked by H3K4me1 or H3K27ac determined by DESeq2 (*P_adj_* < 0.05, |FC| ≥ 1.5). **(C)** Pie charts showing the proportion of accessible enhancers (n = 79056) that represent active enhancers (H3K4me1 + H3K27ac peaks, n = 34438) and proportion of dynamic H3K27ac regions (n = 5100) within active enhancers. **(D)** Motif analysis in regions of lost or gained ATAC-seq accessibility, and regions with dynamic H3K27ac signal in iPB versus niPB. **(E)** Correlation between chromatin accessibility and H3K27ac signal across chromatin sites. Each dot represents a genomic region, with up-regulated sites (yellow) exhibiting increased chromatin accessibility and H3K27ac levels, and down-regulated sites (purple) showing decreased chromatin accessibility and H3K27ac deposition. Red dashed line represents linear regression trend. **(F)** Semantic clustering of GO biological process terms associated with H3K27ac regions exhibiting gained accessibility in iPB relative to niPB. GO terms were identified by GREAT analysis and clustered using the binary-cut algorithm. **(G)** Heatmap of transcription factor target gene modules highlighting upregulation of STAT1 target genes identified by RNA-seq in inflamed versus non- inflamed samples.

Genomic annotation revealed that differential H3K27ac regions were predominantly located within promoter-associated regulatory elements, closely mirroring the distribution observed in the ATAC-seq dataset (fig. S7C). AP-1 family motifs, including FOS, FRA1, ATF3, JUNB, and FRA2, were enriched within regions exhibiting altered accessibility and H3K27ac occupancy (Fig. 4D, and fig. S7, D and F), linking enhancer remodeling to AP-1-associated regulatory programs. Integration of chromatin accessibility and histone modification datasets revealed a strong positive correlation between changes in ATAC-seq accessibility and H3K27ac occupancy, whereas H3K4me1 changes showed minimal association with chromatin accessibility (Fig. 4E). Dynamically regulated enhancer regions were associated with epithelial differentiation, morphogenesis, responses to external stimuli, transport, and apoptosis pathways (Fig. 4F, and fig. S7, F and G), consistent with the altered epithelial differentiation and increased apoptosis observed in iPB organoids. Also, analysis of the RNA-seq dataset showed that STAT1 target genes associated with apoptosis, immune and stress responses, and differentiation programs modules were overexpressed in iPB ISCs (Fig. 4G). Together, these findings demonstrated that the enhancer landscape of pouch body ISCs is impacted by inflammatory status primarily through H3K27ac-dependent remodeling of active enhancers and implicate both AP-1-associated epithelial regulatory programs and STAT1-associated inflammatory programs as major targets of this process.

### Loss of c-Jun alters ISC fate

Because AP-1 motif-containing regulatory regions were extensively remodeled and AP-1/JUN- associated motif accessibility was reduced in iPB ISCs, we investigated whether reduced AP-1 activity contributes to epithelial dysfunction. We observed shallower footprints and lesser chromatin accessibility surrounding BATF::JUN and FOS::JUN motifs indicative of reduced AP-1 occupancy in iPB compared with niPB ISCs in the ATAC-Seq dataset (Fig. 5A). Consistent with these observations, WB analysis demonstrated reduced levels of total and phosphorylated c-Jun in iPB organoids, whereas NF-κB- and MAPK-associated signaling proteins p65, IKKα, and SAPK/JNK were unchanged (Fig. 5, B and C). RT-qPCR confirmed reduced expression of *JUN* and *FOS* in iPB samples (Fig. 5D). Co- immunoprecipitation experiments confirmed that c-Jun formed complexes with AP-1 family members JUNB, BATF, Fra1, and β-catenin (Fig. 5E). c-Jun functions as a scaffold in the β-catenin–TCF complex within the canonical WNT signaling pathway (*6*). Genome browser tracks showed altered chromatin accessibility at regulatory regions associated with *CTNNB1*, *WNT10A*, and *NOTCH1* in ISCs, overlapping with conserved JUN-binding sites identified from publicly available ChIP-seq datasets (Fig. 5F). *JUN* knockdown in niPB organoids by shRNA (Fig. 5E) led to an increase in Annexin V^+^ ISCs and reduced organoid formation efficiency compared to controls (Fig. 5, G to J). *JUN* knockdown also resulted in a decrease in APOA1^+^ cells and increases in MUC2^+^ and CHGA^+^ cells (Fig. 5, K and L). Therefore, loss of AP-1 activity, particularly reduced c-Jun function, was sufficient to increase ISC apoptosis and lineage remodeling.

**Fig. 5.**
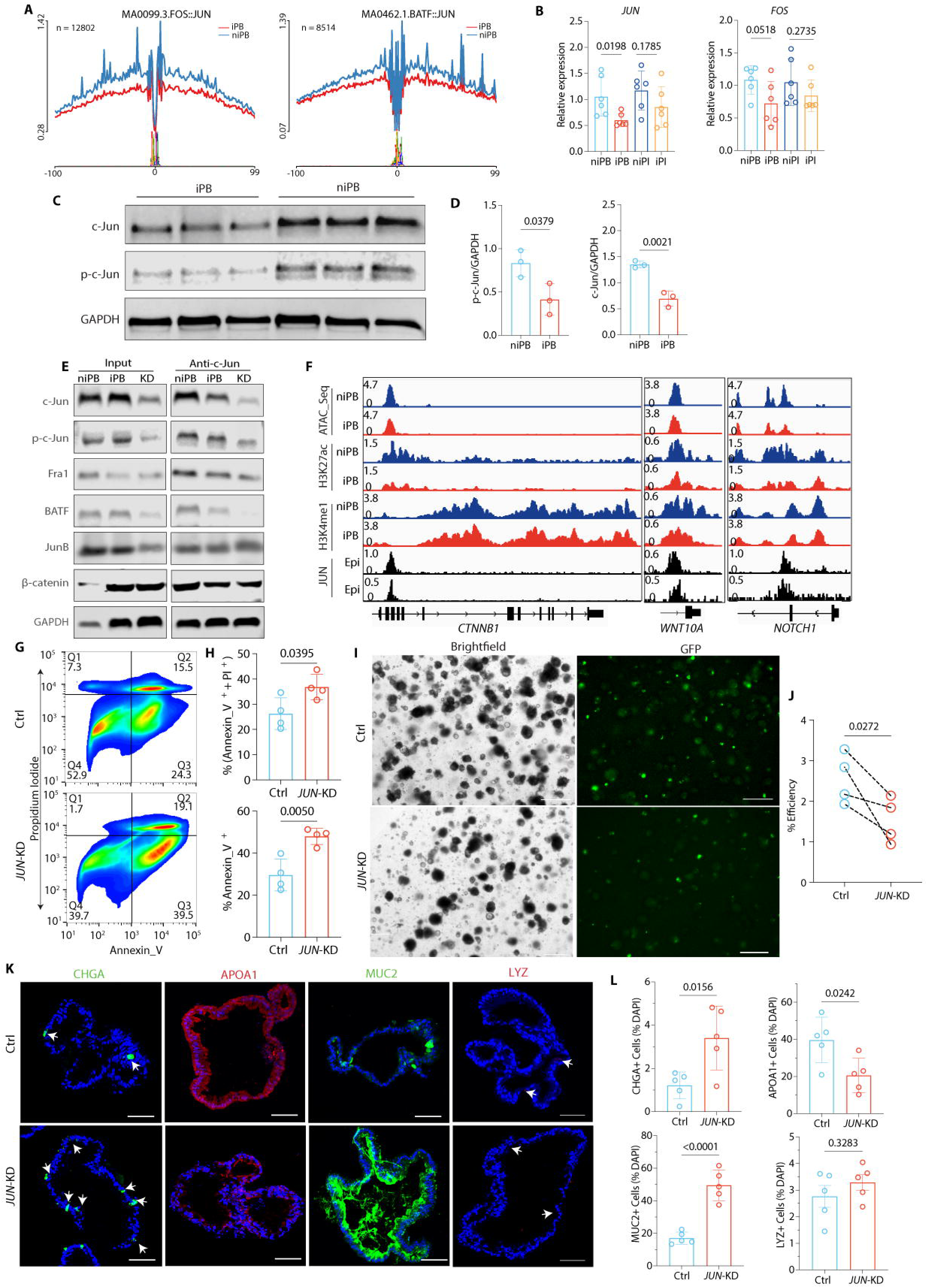
Loss of c-Jun Alters ISC Fate. **(A)** ATAC-seq footprint for BATF::JUN and FOS::JUN motifs in iPB and niPB organoids. Signals across genome motif sites were aligned and averaged. **(B)** RT-qPCR analysis of *JUN* and *FOS* expression in niPB, iPB, niPI and iPI organoids. **(C-D)** WB analysis of c-Jun and p-c-Jun proteins in iPB and niPB organoids (C) and quantification (D). (**E)** Co-immunoprecipitation using anti-c-Jun antibody showing interactions with AP-1 family members in niPB, iPB, and JUN-KD organoids. **(F)** ATAC-seq tracks at the *WNT10A*, *CTNNB1* and *NOTCH1* with public *JUN* ChIP-seq tracks from ChIP-Atlas (ID: SRX10976448; SRX10976239). **(G and H)** Flow cytometry plots (G) and quantification (H) of annexin V and PI stained ISCs in control and *JUN*-KD organoids. **(I and J)** Representative images (I) and quantification of generation efficiency of control and *JUN*-KD (J). GFP indicates shRNA transduction. Scale bars, 400 μm. **(K and L)** Immunofluorescence staining (K) and quantification (L) of shRNA control and *JUN*-KD niPB organoids for CHGA, APOA1, MUC2 and LYZ. Scale bars, 50 µm. **Dots represent individual sample**; bars show the means ± SD. Statistical significance were determined by one-way ANOVA with Tukey’s multiple-comparisons test (B) and paired or unpaired two-tailed Student’s *t*-tests (D, H, J and L). Exact *P* values are shown.

### Pouchitis-associated STAT1 activation impairs ISC function and is reversed by JAK inhibition

To explore strategies for reversing ISC dysfunction, we next tested whether inhibition of upstream JAK signaling could mitigate the STAT1-associated epithelial defects observed in iPB organoids. WB analysis confirmed that STAT1 was phosphorylated (pSTAT1) in iPB organoids (Fig. 6A). We next tested whether inflammatory cytokine TNF-α, IFN-γ, or IFN-λ2 is toxic to pouch organoids. Among these cytokines, IFN-γ produced the strongest reduction in organoid viability in both niPB and iPB cultures (Fig. S8A). Consistent with enhanced pathway activation in vivo, STAT1-positive epithelial cells were significantly increased in inflamed pouch tissues compared with non-inflamed controls (fig. S8, B and C). iPB organoids showed higher expression of several interferon-responsive genes post-differentiation, and this difference at baseline was exaggerated upon stimulation; IFN-γ treatment resulted in a higher fold change in these transcripts in iPB compared with niPB organoids (Fig. 6B, and fig. S8, D and E). Given these findings, we tested whether inhibition of upstream JAK signaling could mitigate inflammation-associated defects. Treatment with tofacitinib, an FDA-approved JAK inhibitor for UC (*34*, *35*), rescued organoid growth and formation efficiency in iPB organoids (Fig. 6C and D). Similar effects were observed with another JAK inhibitor, upadacitinib (fig. S8, F and G). Also, tofacitinib treatment normalized epithelial differentiation of iPB organoids, reversing the expansion of MUC2 cells and reduction of APOA1 cells (Fig. 6, F and G). Overall, these data demonstrated that *STAT1* overexpression associated with inflammation disrupts ISC viability and differentiation.

**Fig. 6.**
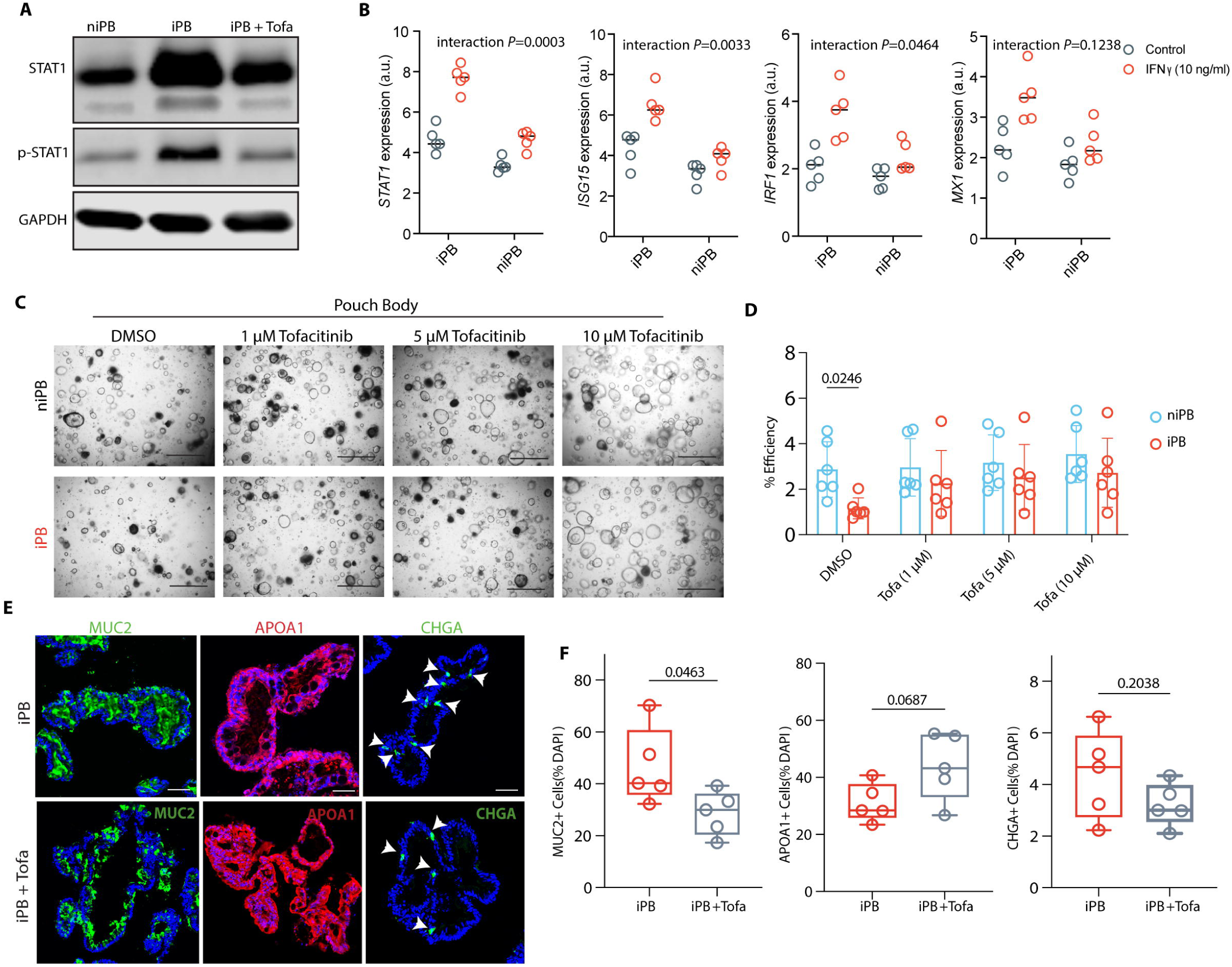
Pouchitis-associated STAT1 activation impairs ISC function and is reversed by JAK inhibition. **(A)** Western blot analysis showing *STAT1* and phosphorylated *STAT1* (p-STAT1) expression in niPB, iPB, and tofacitinib treated inflamed (iPB + Tofa) organoids. **(B)** Quantitative real-time PCR (RT-qPCR) analysis of *STAT1*, *ISG15, IRF1* and *MX1* expression in iPB and niPB organoids following treatment with vehicle control or IFN-γ (10 ng/ml). **(C and D)** Representative brightfield images (C) and quantification of organoid formation efficiency (D) in niPB and iPB organoids treated with the tofacitinib (1, 5, 10 μM). Scale bars, 400 μm. **(E and F)** Immunofluorescence staining (E) and quantification (F) of MUC2, APOA1 and CHGA in iPB and tofacitinib treated iPB organoids. Scale bars, 200 µm. Dots represent individual samples; bars show mean ± SD. For (B), gene expression was analyzed using a linear mixed-effects model (LMM) accounting for donor matching, with gene, condition, treatment, and their interactions as fixed effects and donor as a random effect. Gene-specific condition-by-treatment interaction P values are shown. Statistical significance determined by two-way ANOVA with Holm–Šidák multiple-comparisons correction (D) and unpaired two-tailed Student’s *t*- tests (F). Exact *P* values are shown.

### Tofacitinib reverses inflammation-associated chromatin remodeling in ISCs

To investigate the impact of JAK/STAT inhibition on chromatin accessibility, we performed ATAC-seq on ISCs isolated from tofacitinib-treated iPB organoids and compared them with untreated iPB and niPB controls. Tofacitinib treatment led to substantial changes in the chromatin landscape of iPB organoids (Fig. 7, A and B). PCA demonstrated that tofacitinib shifted the chromatin landscape of iPB towards the non-inflamed state along the PC1 axis (fig. S8H). Consistently, direct comparison of tofacitinib-treated iPB with niPB showed fewer differentially accessible regions, further indicating that JAK inhibition shifted the chromatin landscape of iPB ISCs toward the non-inflamed state (fig. S8I). Among the 7,508 regions that gained accessibility in iPB relative to niPB, 57.4% reverted to a closed state following tofacitinib treatment (Fig. 7C). Tofacitinib-rescued regions were predominantly located in promoter and intronic regions, whereas the remaining 42.6% were classified as Tofa-resistant, indicating accessibility changes that persisted despite tofacitinib treatment (Fig. 7D). Regions losing accessibility after treatment were highly enriched for AP-1 family motifs, whereas newly accessible regions were enriched for GATA-family TF motifs associated with epithelial differentiation programs (Fig. 7E). Chromatin regions altered by tofacitinib are in loci associated with epithelial development and differentiation, cellular responses to external stimuli, leukocyte-associated signaling, and transport processes (Fig. 7F). Accessibility at epithelial regulatory genes including *HES1*, *TGFB2*, *EGR3*, *CTNNB1*, *CASP7*, and *ATOH1* was remodeled following treatment, and *NEUROD3* expression was reduced alongside increased *HES1* expression (Fig. 7, G and H). These findings suggested that JAK inhibition promotes re-establishment of epithelial regulatory networks while suppressing inflammation-associated chromatin states.

**Figure 7.**
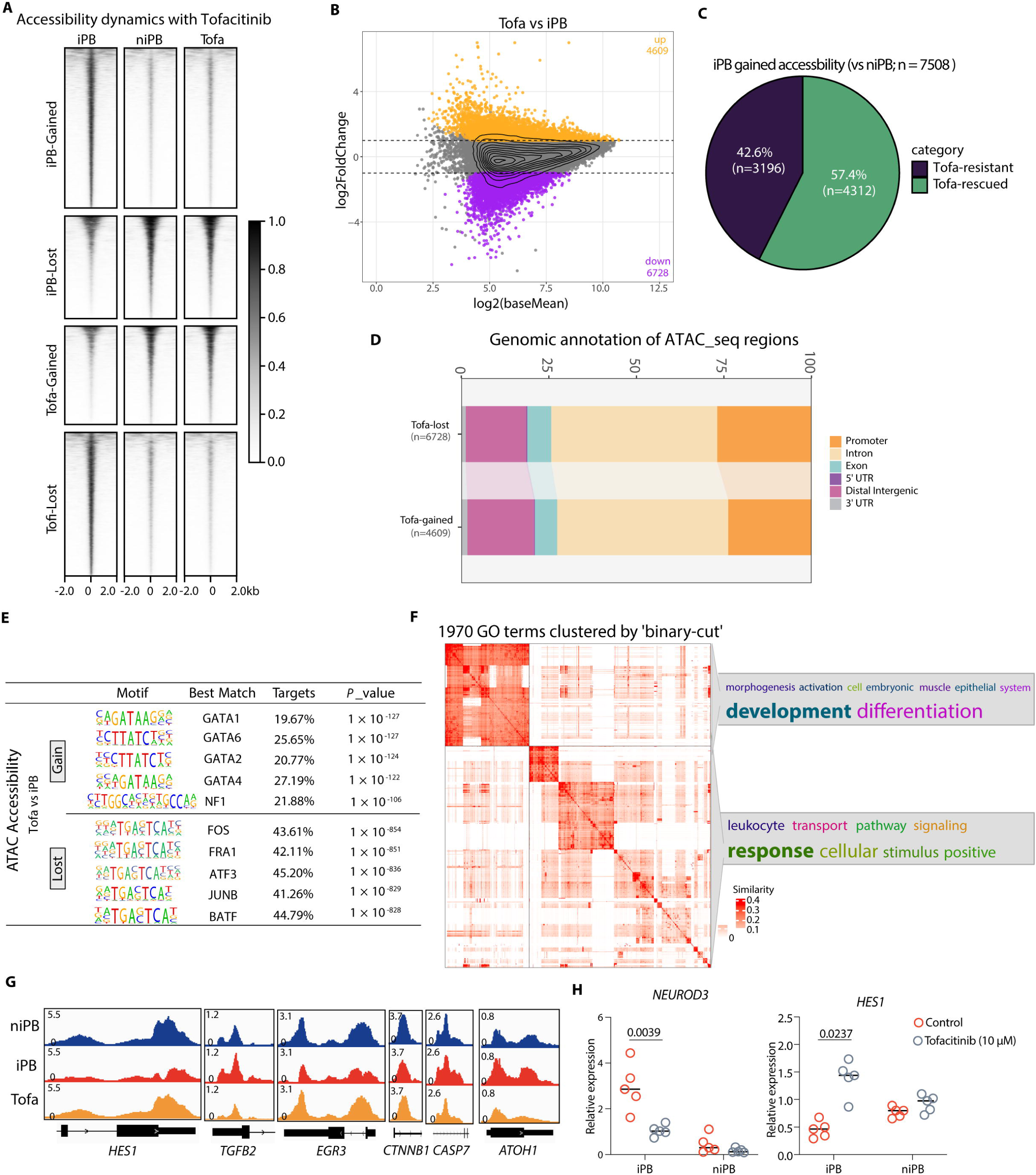
Tofacitinib reverses inflammation-associated chromatin remodeling in pouch ISCs. **(A)** Heatmaps of ATAC-seq accessibility in niPB, iPB, and tofacitinib-treated iPB (Tofa) ISCs across differentially accessible regions, centered on peak summits (±2 kb). **(B)** MA plot showing regions of lost and gained chromatin accessibility in Tofa compared with iPB. Significant increase or decrease was determined by DESeq2 (*P_adj_* < 0.05, |FC| ≥ 1.5). **(C)** Proportion of inflammation-induced accessible regions in iPB that were closed following tofacitinib treatment (Tofa-closed) versus regions that remained accessible. **(D)** Genomic annotation of regions gained or lost following tofacitinib treatment. (E) Motif enrichment analysis of regions differentially regulated by tofacitinib treatment. **(F)** Gene Ontology (GO) biological process terms associated with chromatin regions altered following tofacitinib treatment. GO terms were identified by GREAT analysis and clustered using the binary-cut algorithm. **(G)** ATAC-seq tracks showing accessibility changes at epithelial regulatory genes, including *HES1, TGFB2, EGR3, CTNNB1, CASP7,* and *ATOH1*, following tofacitinib treatment. **(H)** RT-qPCR analysis of *NEUROD3* and *HES1* expression in niPB and iPB organoids with or without tofacitinib treatment. Dots represent individual samples; horizontal lines indicate mean. Statistical significances were determined by one-way ANOVA with Tukey’s post hoc test. Exact P values are shown.

## DISCUSSION

We found that de novo inflammation in a surgically generated intestinal organ, the J-pouch, leads to persistent damage to the ISC compartment, as evidenced by reduced organoid formation and increased apoptosis after multiple rounds of passaging. Additionally, organoids derived from an inflamed J-pouch displayed increases in enteroendocrine and goblet cells and reduction in enterocytes, which was also observed in primary tissue. These defects in cell death and differentiation were associated with major disruptions to the chromatin landscape in ISCs, primarily through histone acetylation changes at active enhancers. Whereas AP-1 TF motifs such as c-Jun binding sites became closed, loci associated with JAK/STAT signaling were hyperactive. Inhibiting c-Jun in organoids to recreate inflammation-induced deficits in AP-1 TF expression and binding site accessibility led to enhanced cell death and secretory cell differentiation. Conversely, inhibiting the overactive JAK/STAT largely reversed ISC defects in pouchitis organoids and restored chromatin accessibility at AP-1 TF motifs, indicating that the aberrant immune signaling mediates chromatin level changes. When taken together, these findings support a model in which excess JAK/STAT activity causes persistent changes in the ISC compartment of inflamed pouch tissue through epigenetic changes associated with reduced c-Jun function.

Eligibility for J-pouch surgery typically requires the absence of ileal inflammation at the time of pouch creation. Therefore, we can be confident that the ISCs analyzed in this model are derived from newly developing inflammation in previously unaffected ileal tissue. By demonstrating long-lasting chromatin level changes in ISCs, our findings help explain prior studies showing that organoids derived from IBD patients differ from non-IBD controls (*14*). Organoids derived from the ileum and colon of patients with IBD have been reported to display growth defects (*13*, *36*), suggesting that propensity for cell death is a general feature of injured ISCs in inflamed tissue. Although upregulation of immune transcripts is another common theme, the expression of epithelial lineage markers in IBD organoids vary considerably between studies. Colonic organoids from pediatric patients with UC display reductions in goblet cells and enhanced glycolysis (*37*). However, single-cell RNA-seq of small intestinal organoids identified stem cell and metabolic pathway markers as distinguishing features when comparing samples from inactive versus active CD rather than secretory cell changes (*38*). Consistent with anatomical region-specificity, the consequence of inflammation was consistently less dramatic on the pouch inlet organoids and tissue in our study. This regional difference may reflect the distinct anatomy and function of the pouch body (*39*), which serves as the primary stool reservoir and is therefore exposed to prolonged contact with microbial communities, luminal metabolites, mechanical stress, and immune stimulation (*40–43*). In contrast, the pouch inlet is anatomically closer to the neo-terminal ileum and may retain a more ileal-like environment and less exposure to fecal stasis. Clinically, this distinction may be important because it suggests that pouchitis-associated epithelial injury may not be uniform throughout the pouch, and that the pouch body may represent a particularly vulnerable site for persistent ISC dysfunction, impaired epithelial repair, and recurrent inflammation.

The AP-1 TF family regulates enhancer dynamics and transcriptional programs essential for maintaining stem cell function under homeostatic and pathological conditions (*44–46*). Our findings provide evidence that AP-1 activity contributes to ISC maintenance in humans. In stark contrast to the murine skin injury model, where increased AP-1 TF activity promotes chromatin accessibility and injury resolution (*15*), inflamed pouch ISCs displayed reduced AP-1 activity, exacerbating ISC loss and dysfunction. AP-1 can both regulate enhancer accessibility and respond to changes in chromatin state. Our data reveal reductions in chromatin accessibility and H3K27ac modifications in ISCs in iPB compared with niPB, suggesting that AP-1 suppression and enhancer remodeling may reinforce one another to destabilize ISCs. Based on our experiments in which we treat organoid with IFN-γ or JAK/STAT inhibitors, it is possible that pro-inflammatory cytokines (*47*, *48*) or microbiome-derived products recruit chromatin remodelers such as BRG1 and SWI/SNF (*49*) to drive enhancer remodeling, leading to c-Jun depletion and a persistently altered ISC fate.

We acknowledge several limitations to this work. First, although patient-derived pouch organoids preserve clinically relevant epithelial features, the sample size is limited by the availability of pouch biopsies, as well as technical and throughput constraints associated with establishing and profiling organoid lines in parallel. Second, organoids allow epithelial-intrinsic mechanisms to be examined but do not fully capture interactions with immune cells, stromal cells, and microbiota that likely contribute to pouchitis in vivo. Third, it is difficult to establish the causal relationship between a regulatory region for a specific locus and functional outcomes. It is more likely that the AP-1 and JAK/STAT axis act on multiple sites in concert rather than through a simple linear manner. Lastly, although tofacitinib restored many of the alterations to chromatin accessibility associated with inflammation, the effect of the drug was incomplete, indicating that other signals contribute to persistent ISC defects. Additional studies will be needed to determine whether restoring AP-1 activity and JAK inhibition can reverse ISC injury in a durable manner.

Disease occurring within the first 180 days after IPAA surgery is a known predictor of future chronic pouchitis (*50*, *51*). The involvement of overactive STAT1 would be consistent with a mechanism in which ISCs in previously inflamed tissue become sensitized to cytokines such as IFN-γ. Although JAK inhibition is well known to dampen cytokine-driven immune cell activation (*52*, *53*), our data raise the possibility that they can be used to suppresses persistent *STAT1* activation in the pouch epithelium. JAK inhibitors have been evaluated clinically in CD and UC, and are FDA-approved for moderate-to-severe disease (*34*, *35*), but their application in pouchitis remains limited (*54*, *55*). In addition to supporting a more careful evaluation of JAK inhibitors for the prevention or treatment of pouchitis, our findings raise the intriguing possibility of using interventions to reset defective epigenetic programs. For instance, short-chain fatty acids and ketone bodies act as HDAC inhibitors to regulate ISC fate and intestinal homeostasis (*56–58*). With a better understanding of the mechanism and consequence of ISC injury, it may be possible to combine traditional immune-targeting medications with those that reverse epigenetic modifications or their signaling consequences to achieve a sustained response in patients with pouchitis and other intestinal disorders.

## MATERIALS AND METHODS

### Study design

This study aimed to investigate how inflammation alters ISC function and contributes to the pathophysiology of pouchitis in the human J-pouch. Human participants were not randomly assigned because study groups were defined by clinical inflammatory status and anatomical location of the biopsy. Participant inclusion and exclusion criteria were established before sample collection. Organoids were generated from all curated samples with matched pouch inlet and body biopsies available at the time of study initiation. Exclusion criteria were missing clinical data or insufficient viable material. Histological assessment of the tissue for determination of inflammatory status was performed blind by pathologists. Patient-derived J-pouch organoids and matched primary tissue sections were used to assess ISC phenotypes affected by pouchitis. ISC survival and organoid-forming capacity were evaluated using organoid culture assays. Proliferation was assessed by Ki-67 immunofluorescence staining, whereas apoptosis was evaluated by flow cytometry and Western blotting for apoptosis-associated markers. Epithelial differentiation was assessed by immunofluorescence staining and RT-qPCR analysis of lineage markers in organoids, with selected findings validated in matched primary tissue sections.

To define regulatory mechanisms associated with inflammation, CD24 CD166 CD44 PTK7^high^ ISCs were isolated by flow cytometry and profiled by ATAC-seq, RNA-seq, and H3K27ac/H3K4me1 CUT&RUN. These approaches were used to assess chromatin accessibility, transcriptional changes, and histone modification patterns in ISCs. Mechanistic intervention experiments were performed using apoptosis inhibitors, JUN knockdown, and inflammatory cytokine stimulation. To test therapeutic reversibility, JAK inhibition experiments were performed using tofacitinib or upadacitinib, with tofacitinib used to determine whether JAK inhibition could restore ISC function and chromatin states altered by inflammation.

Unless otherwise indicated, experiments were performed using at least three biologically independent patient-derived samples or organoid lines. Technical replicates, when used, were averaged within each biological replicate and were not treated as independent observations. Sample sizes were based on our prior related studies, similar published studies, availability of human tissue and successful establishment of organoid lines. No statistical method was used to predetermine sample size. No data points were excluded as statistical outliers. Unless otherwise indicated, experiments were performed using biologically independent tissue or patient-derived samples.

### Human specimens and tissue collection

Patients with non-IBD, UC and CD were recruited at outpatient colonoscopy performed for colon cancer screening, surveillance, or IBD activity assessment at New York University (NYU) Langone Health’s Ambulatory Care Center, New York, under an NYU Grossman School of Medicine Institutional Review Board–approved study (Mucosal Immune Profiling in Patients with endoscopy from the Inflammatory Bowel Disease; S12-01137). Mucosal pinch biopsies were obtained via pouchoscopy from the ascending colon of UC patients with a J-pouch approximately six months to several years after final stage of IPAA surgery, when continuity and continence were restored. In 20 consented J-pouch patients, biopsies were obtained from the J-pouch body and inlet, and subsequently stratified by inflammation status via visible endoscopic appearance. Inflammation status was also confirmed by endoscopic and expert IBD histopathological examination. Potential participants were excluded if clinical presentation was consistent with “Crohn’s disease of the pouch”, defined by the presence of fistula/fistulae, stricture involving the pouch or pre-pouch ileum, and the presence of long-segment pre-pouch ileitis, or if they did not provide tissue.

### Statistical analysis

Individual-level tabular data are presented in data file S1. Statistical analyses were performed using GraphPad Prism and R software. Unless otherwise indicated, data are presented as mean ± SD. Comparisons between two groups were made using paired and unpaired two-tailed Student’s t-tests, as appropriate. For comparisons among more than two groups, one-way ANOVA followed by Tukey’s post hoc test was used. For experiments involving two independent variables, two-way ANOVA followed by Holm–Šidák correction for multiple comparisons was used. Correlations were assessed using the Pearson or Spearman correlation according to data distribution. For the IFN-γ recall experiment, gene expression was analyzed using a linear mixed-effects model in R, with gene, condition, treatment, and their interactions as fixed effects and donor as a random effect. Comparisons of ATAC-seq, CUT&RUN, and RNA-seq signals between two groups were performed using Welch’s two-sample t test, and differences in cumulative distributions were assessed using the Wilcoxon test. All statistical tests were two-sided unless otherwise indicated, and an alpha level of 0.05 was used.

### List of Supplementary Materials

Materials and Methods

Fig. S1 to S8

Table S1 to S4

References (*59–77*)

MDAR Reproducibility Checklist

Data file S1

## Acknowledgments

We would like to acknowledge NYU Grossman School of Medicine Reagent Preparation Core, Microscopy Laboratory, Genome Technology Center, and Center for Biospecimen Research Development for use of their instruments and technical assistance. We also acknowledge the Penn medicine of Cell & Developmental Biology (CDB) Microscopy Core (RRID SCR_022373), Penn Genomics and Sequencing Core, Penn Molecular Pathology & Imaging Core (RRID: SCR_022420) funded by the Center for Molecular Studies in Digestive and Liver Diseases (NIH P30 DK050306) and CHOP Flow Cytometry Core for their assistance and resources.

## Author contributions

K.C., C.Z., and K.K.J. conceived the study and designed the experiments. C.Z., T.L., S.L., and K.K.J. performed and analyzed the organoid culture, Western blotting, and RT-qPCR experiments. C.Z. performed flow cytometry and FACS-based assays, human tissue immunofluorescence, co-immunoprecipitation, and functional perturbation experiments. C.Z., S.L., and K.K.J. performed sequencing library preparation. C.Z. analyzed the RNA-seq, ATAC-seq, and CUT&RUN data. K.K.J. and K.S.V. generated the human organoids. C.Z. and J.Y.C.C.F. assisted with human biopsy collection. J.E.A. and K.C. supervised the study. J.E.A., K.K.J., and K.C. acquired funding. C.Z. and K.C. wrote the original draft of the manuscript. All authors reviewed and edited the manuscript and approved the final version.

## Data, code, and materials availability

All data associated with this study are in the paper or supplementary materials. Genome data used in this study have been deposited with the National Center for Biotechnology Information Gene Expression Omnibus (GEO) under accession number GSE338895. Analysis code used in this study is available at Zenodo (DOI: 10.5281/zenodo.22073650). Human biopsy specimens and patient-derived organoid lines are subject to the terms of the approved IRB protocol and applicable institutional requirements and may require institutional approval before sharing. All other materials used or generated in this study are commercially available or will be supplied upon reasonable request.

