## Supplementary Materials and Methods, Figures, and Tables for "Epigenetic Reprogramming Alters Human Intestinal Stem Cell Fate in Pouchitis"

**Culture of human organoids**

We adapted a described protocol for the generation and expansion of J-pouch organoids (*1*, *2*). Biopsies were collected in ice-cold complete Roswell Park Memorial Institute (RPMI) medium [RPMI 1640 supplemented with 10% fetal bovine serum (FBS), 100 IU Penicillin and 100 μg/ml Streptomycin (Corning), 2 mM L-Glutamine (Corning), and 50 μM 2-mercaptoethanol (ThermoFisher)] and incubated in Gentle Cell Dissociation Reagent (Stem Cell Technologies) on ice for 30 minutes followed by vigorous pipetting to isolate crypts. Crypts were embedded in 30 μl of Matrigel and cultured with Human IntestiCult Organoid Growth Medium (Basal Medium and Organoid Supplement, Stem Cell Technologies). The culture medium was changed every 3 days. Organoids were passaged once a week and 10 μM Y-27632 was added for the first 3 days.

**Organoid differentiation, formation efficiency, and viability assays**

Organoids were harvested using Cell Recovery Medium (Corning) and incubated at 37°C for 15 minutes to generate single cell suspensions using TrypLE Express enzyme (ThermoFisher), and a uniform suspension containing the same number of cells for each sample was prepared in Matrigel. For differentiation, the maintenance medium was replaced with DMEM/F-12 (ThermoFisher) supplemented with 100 IU/mL Penicillin, 100 μg/mL Streptomycin, 125 μg/mL Gentamicin (ThermoFisher), 2 mM L-Glutamine, 20 ng/mL recombinant epidermal growth factor (EGF) (PeproTech), 100 ng/mL Noggin (R&D Systems), 1 μg/mL R-Spondin 1 (R&D Systems), 100 ng/mL recombinant Wnt-3a (R&D Systems), 1 mM N-acetylcysteine (Sigma-Aldrich), 10 nM Gastrin (Sigma-Aldrich), 500 nM A-83-01 (Tocris), 200 ng/mL insulin-like growth factor 1 (IGF-1) (BioLegend), 100 ng/mL fibroblast growth factor 2 (FGF-2) (PeproTech), and 1x B27, referred to as IF medium (IGF-1 and FGF-2 medium). Each 10 μl droplet of Matrigel containing 3000 cells were cultured in a 96-well culture plate in triplicate, supplemented with 10 μM Y-27632 for the first three days to promote cell survival. To assess organoid formation efficiency, the single cells plated in Matrigel (Corning) in 96-well plates at a density of 3000–6000 cells per well, cultured in differentiation media. Organoid formation was evaluated by counting the number of formed organoids under a brightfield microscope at 7–10 days post-plating. Formation efficiency was calculated using the formula: Efficiency (%) = (Number of formed organoids / Number of seeded cells) × 100. For thiazolyl blue tetrazolium bromide (MTT) reduction assay, staining with MTT was adapted from a previously described method (*1*). In brief, we added 10 µl MTT (Sigma-Aldrich; 5mg/mL) into the organoids. After incubation for 2 h at 37°C, 5% CO2, the medium was discarded and 20 μl of 2% SDS (Sigma-Aldrich) solution in water was added to solubilize the Matrigel for 2 hrs. Then, 100 μl of DMSO (ThermoFisher) was added for 1 h to solubilize the reduced MTT, and OD was measured on a microplate absorbance reader (ParkinElmer) at 562 nm. The specific organoid death (%) was calculated as MTT reduction (%) by normalizing to untreated organoids which were defined as 100% viable. To assess the effects of chemical inhibitors, organoids were co-cultured for 5 to 7 days with the following: 1 μM Q-VD-OPh (pan-caspase inhibitor, Sigma-Aldrich), 10 μM Z-DEVD-FMK (caspase-3 inhibitor, MedChemExpress), 20 μM Z-IETD-FMK (caspase-8 inhibitor, MedChemExpress), 10 μM Tauroursodeoxycholic Acid (TUDCA, Selleckchem), and 10 μM Z-LEHD-FMK TFA (caspase-9 inhibitor, Selleckchem), Tofacitinib (JAK inhibitor, Sigma-Aldrich), Upadacitinib (JAK inhibitor, Sigma-Aldrich).

**Flow cytometry**

The following antibodies were used for stem cell sorting: CD326 (565685, 1:50) from BD Biosciences, CD44 (103040, 1:200) and CD24 (311116, 1:200) from Biolegend, CD166 (12-1668-421, 1:200) from eBioscience, and PTK7 (130-091-366, 1:50) from Miltenyi Biotec. Apoptosis was assessed using the Annexin V-FITC Apoptosis Detection Kit (BD Biosciences) (*3*, *4*). Organoids were collected using Cell Recovery Solution (Corning, 354270) and digested into single cells with TrypLE Express Enzyme (Gibco, 122604021) for 15 minutes in a 37 °C water bath. The resulting cells were stained with stem cell markers for 20 minutes at 4 °C in FACS buffer (PBS supplemented with 2% FCS, 10 µM Y27632, 2 mM EDTA, and 1 mM NAC), washed, and resuspended in the same buffer. DAPI (ThermoFisher) was used to exclude dead cells during stem cell sorting. Apoptosis staining was performed according to the manufacturer’s protocol. Briefly, cells were washed twice with cold PBS and resuspended in binding buffer. Cells were stained with Annexin V-FITC and propidium iodide for 15 minutes at room temperature in the dark. Stained cells were analyzed using a Cytek Aurora flow cytometer analyzer (Cytek Biosciences), and data were processed using FlowJo software. Apoptosis was assessed using the Annexin V-FITC Apoptosis Detection Kit (BD Biosciences). Organoids were collected using Cell Recovery Solution (Corning, 354270) and digested into single cells with TrypLE Express Enzyme (Gibco, 122604021) for 15 minutes in a 37 °C water bath. The resulting cells were stained with stem cell markers for 20 minutes at 4 °C in FACS buffer (PBS supplemented with 2% FCS, 10 µM Y27632, 2 mM EDTA, and 1 mM NAC), washed, and resuspended in the same buffer. DAPI was used to exclude dead cells during stem cell sorting.

**ATAC Sequencing**

ATAC-seq was performed as previously described (*5*, *6*). 50k~100k FACS-sorted stem cells, gated on CD24^+^CD166^+^CD44^+^PTK7^high^, were collected and pelleted by centrifugation at 500 *x* *g* at 4°C for 5 mins. Nuclei were exacted in ATAC-Resuspension Buffer and incubated with Tn5 transposase at 37°C for 30 min. Size distribution of the amplified DNA was analyzed using High-sensitivity Qubit dsDNA Assay Kit (ThermoFisher). Libraries were pooled for paired-read sequencing performed on NovaSeq 6000 (Illumina) at the NYU Genome Technology Core.

ATAC-seq reads underwent quality control and adapter trimming using FastQC software. The reads were then aligned to the GRCh38/hg38 reference genome using Bowtie2 (*7*). Peaks were identified using Genrich in ATAC-seq mode, applying a q-value filter of < 0.01 to ensure the reliability of peak calls. To visualize the data, normalized fragment signal bigWig files were created using DeepTools2 (*8*). The normalization method employed was BPM (Bins Per Million), ensuring that the data was comparable across samples. The bigWig files were generated with a bin size of 10 bases and were centered on the reads' fragment length. Additionally, normalized signals ± 2 kb from the center of the ATAC-seq peaks were utilized for unsupervised k-means clustering analysis, also utilizing DeepTools for this step. A list of ATAC-seq consensus peak set was made using DiffBind (*9*). Differential enrichment was analyzed using DESeq2 (*10*) with significant changes being defined by FDR value <0.05 and |FC| ≥1.5 (gain or loss). Ontology analysis was performed using Genomic Regions Enrichment of Annotation Tool (GREAT) (*11*) with the human genome (GRCh38/hg38) as the background. For the identification of TF footprints in ATAC-seq peaks, we used the HINT tool of the Regulatory Genomics Toolbox (*12*). Motifs enriched in ATAC regions were identified using the HOMER *de novo* algorithm (*13*) based on the cumulative binomial distribution and the annotates peaks function was used. To characterize the distribution of binding sites in ATAC-seq data, peak sites were mapped to various annotations, including transcription start site (TSS), transcription termination site (TTS), exon (coding), 5’ UTR exon, 3’ UTR exon, intronic, and intergenic regions. These annotations are commonly defined by HOMER.

**CUT&RUN assay**

ISCs were sorted by FACS based on CD24^+^CD166^+^CD44^+^PTK7^high^ surface markers for and processed using the CUT&RUN Assay Kit (Cell Signaling Technologies, #86652) according to the manufacturer's instructions. The cell suspension was first incubated with concanavalin A beads, then with 4 µg Acetyl-Histone H3 (Lys27) (H3K27ac, Cell Signaling Technologies, #8173S) or Mono-Methyl-Histone H3 (Lys4) (H3K4me1, Cell Signaling Technologies, #5326S) antibodies at 4°C for 1 hr. Chromatin-bound beads were mixed with pAG-MNase in digitonin buffer, and MNase activity was activated by adding cold calcium chloride, followed by incubation at 4°C for 30 minutes. De-crosslinking was achieved through sequential treatment with RNase at 37°C for 10 minutes and proteinase K at 65°C for 2 hrs. Enriched DNA was purified using the DNA Purification Kit (Cell Signaling Technologies, #14209S) and prepared for sequencing using the DNA Library Prep Kit for Illumina (Cell Signaling Technologies, #56795). Library quality and size distribution were assessed using the Bioanalyzer High Sensitivity DNA Analysis Kit (Agilent, #5067-4626) according to the manufacturer’s supplied protocol, and libraries were sequenced as paired-end 100 bp reads (PE100) on a NovaSeq 6000 platform (Illumina) at the Penn Genomics and Sequencing Core.

The FastQC files were processed similar to ATAC-seq, including the procedures of trimming, mapping, and filtering. For peak calling, H3K27ac and H3K4me1 peaks were called using MACS2 (*14*) against input sample of each cell type with a P value cutoff of 1 × 10^−9^ and were filtered against hg38 ENCODE blacklisted region. A list of H3K27ac consensus peak set was made using DiffBind. Accessible enhancers that overlap with any H3K27ac peaks were defined as active enhancers. Differential enrichment of chromatin marks (H3K4me1 and H3K27ac) was analyzed using DESeq2, with significant changes being defined by false discovery rate (FDR) value < 0.05 and |FC| ≥ 1.5 (gain or loss). Accessible enhancers with gain or loss of H3K4me1 levels between conditions were defined as dynamic enhancers; likewise, those with gain or loss of H3K27ac were considered as having dynamic activity.

**RNA-sequencing**

Total RNA was extracted from sorted ISCs using RNeasy Mini Kit with DNase treatment (Qiagen). The libraries for RNA-Seq analysis were generated using TruSeq Standard Total RNA Library Prep with RiboZero Gold rRNA removal kit (Illumina) according to the manufacturer’s supplied protocol. Sequencing was performed on NovaSeq 6000 (Illumina) at NYU Genome Technology Core.

Raw sequencing FastQC files were assessed for quality, adapter content and duplication rates with FastQC. Raw reads were aligned to the human genome (GRCh38 assembly) using HISAT2 (*15*). Transcript levels were quantified as read counts with HTSeq (*16*) . Data normalization and sample variability were assessed through principal component analysis (PCA) using **DESeq2**. Differential expression analysis was performed in **DESeq2,** with thresholds for significance set at an |FC| > 1.5 and a false discovery rate (FDR) < 0.05. Gene set enrichment analysis was performed using GSEA software (Broad Institute) to identify pathways significantly enriched in inflamed versus non-inflamed conditions. RNA-Seq data were used to rank genes based on their differential expression, and the enrichment of specific gene sets was evaluated using the KEGG and GO biological processes databases.

**Western blotting and immunoprecipitations**

J pouch organoids were washed with PBS, incubated with Cell Recovery Solution (Corning) at 4°C for 60 min to dissociate Matrigel, and centrifuged at 400 *x g* for 5 min. The pellets were suspended in RIPA lysis buffer (Sigma-Aldrich) with 2× Halt Protease and Phosphatase Inhibitor Cocktail (ThermoFisher)] and pelleted at 10,000 *x g* for 15 min at 4°C to collect the lysates. The organoid lysates were resolved on Bolt 4-12% Bis-Tris Plus Gels (Invitrogen), transferred onto polyvinylidene difluoride membranes, and blocked using Intercept (TBS) blocking buffer (LI-COR). Membranes were probed with primary antibody overnight at 4°C. The following primary antibodies used were purchased from Cell Signaling except for β-actin (Sigma-Aldrich, A5441): anti-PARP (9542S), caspase-8 (4790S), caspase-9 (9508S), caspase-7 (9494S), caspase-3 (9662S), c-JUN (9165S), Phospho-c-JUN (Tyr701) (3720S), FOS (74620S), BATF(D7C5), JUNB (3758S), β-Catenin (12475S), GAPDH (2118S) and NF-κB Pathway Antibody Sampler Kit (9936). After incubation with the primary antibody, the membrane was washed and probed with the secondary antibody for 1 hr at room temperature. As for secondary antibodies, IRDye 680RD Goat anti-Rabbit (925-68071) and IRDye 800CW Goat anti-Mouse (925-32210) were purchased from LI-COR. After additional washing, protein was detected with Image Studio for Odyssey CLx (LI-COR). Band intensities were measured by Fiji/ImageJ.

Immunoprecipitations were performed with 100 μg whole organoid protein lysates harvested by nondenaturing NP-40 lysis buffer (ThermoFisher) and protein concentrations were estimated by Bio-Rad colorimetric assay. This protein extract was then mixed with Dynabeads Protein G (Invitrogen) that had been pre-coupled with c-Jun (9165S) antibody. This mixture was incubated at 4°C for overnight. After the incubation, the supernatant was removed, and the beads were washed twice with lysis buffer. The beads were subsequently boiled in SDS buffer at 100°C for 3 min and fractionated on Bolt 4-12% Bis-Tris Plus Gels. Detection of immunoprecipitated proteins was performed with above mentioned reagents and antibodies.

**RNA isolation and RT–qPCR**

Total RNA was extracted from dissociated organoids using the RNeasy Mini Kit (Qiagen, 74104) according to the manufacturer’s protocol. Complementary DNA (cDNA) was synthesized using the High-Capacity cDNA Reverse Transcription Kit (Fisher Scientific, 4368813). Quantitative real-time PCR (RT-qPCR) was conducted on a QuantStudio 5 Real-Time PCR System using SYBR Green Master Mix (ThermoFisher, A46111) and gene-specific primers. Gene expression levels were normalized to β-actin as an internal control. Relative gene expression levels were calculated using the ΔΔC_T_ method (*17*).

**Immunofluorescence**

Mucosal pinch biopsies of the ascending colon, J-pouch, and ileum were fixed in 10% formalin and embedded in paraffin blocks. Sections were cut to 5 μm thickness at the NYU Center for Biospecimen Research and Development and mounted on frosted glass slides. For deparaffinization, immersing slides in xylene 15 mins and 1:1 xylene:100% ethanol for 5 minutes, followed by rehydration in descending concentration of ethanol (100%, 95%, 80%, 70%, 50%, ddH_2_O twice; each step for 5mins). Heat-induced epitope retrieval was performed in the antigen retriever 2100 (Electron Microscopy Sciences) pressure cooker using R-UNIVERSAL Epitope Recovery Buffer (Electron Microscopy Sciences) follow the cooker set process. After further cooling to room temperature Tissue areas were circled using a PAP pen, followed by the addition of a permeabilization buffer (PBS containing 0.1% Tween-20 and 0.01% Triton X-100) for 20 minutes at room temperature in humidified chamber. The permeabilization buffer was discarded without washing. Sections were then blocked with BlockAid™ Blocking Solution (ThermoFisher) for 1 hr at room temperature, followed by incubation with primary antibodies anti-APOA1 (rabbit, 1:500, ThermoFisher, PA5-88109), anti-MUC2 (rabbit, 1:500, Santa Cruz, SC-15334), anti-CHGA (mouse, 1:100, Santa Cruz, SC-393941), anti-LYZ (rabbit, 1:500, Abcam), anti-OLMF4 (rabbit, 1:200, Cell signaling,) diluted in blocking buffer. Slides were incubated overnight at 4°C in a humidified chamber. On the following day, sections were washed three times for 10 minutes each in PBS. Secondary antibodies 1:500 diluted in blocking buffer were applied for 1 hr at room temperature. Slides were washed three times for 5 minutes each in PBS and incubated with DAPI (1:1000, ThermoFisher) diluted in PBS for 10 minutes at room temperature. Following this, slides were washed again three times for 5 minutes in PBS. Slides were dried at room temperature for 30 minutes and mounted with ProLong Gold Antifade Mountant buffer (ThermoFisher). Slides were allowed to dry overnight at room temperature before sealing with coverslip sealant. Stained sections were stored at 4°C. Images were captured using Zeiss LSM 980 confocal microscope and analyzed using Fiji.

For human intestinal organoids, frozen sections were prepared as previously described (*18*). Briefly, differentiated organoids were fixed in 4% paraformaldehyde (Electron Microscopy Sciences) and cryoprotected with 20% sucrose (Sigma-Aldrich). Fixed organoids were embedded in NEG-50 (Epredia) and frozen in cryomolds using 2-methylbutane (Sigma-Aldrich) cooled with dry ice. Sections were cut to a thickness of 10 µm using a Cryostat (Micron HM350; ThermoFisher). Staining was performed as described above.

**Lentivirus infection and gene knockdown**

Lentiviral infection and *JUN* gene knockdown in organoids were performed as previously described (*1*, *19*). Briefly, the c-JUN (JUN) Human shRNA Plasmid Kit (TL320397) and a scrambled shRNA control in the pGFP-C-shLenti shRNA Vector (TR30021) were purchased from Origene. Each lentiviral construct, along with Lentiviral Packaging Kits (TR30037), was co-transfected into 293FT cells following the manufacturer’s protocol. The supernatant containing lentivirus was collected and concentrated using the Lenti-X concentrator (Clontech). Organoids derived from the J-pouch were cultured as described above. On day 5, the organoids were mechanically dissociated into single cells by gentle pipetting and incubated with TrypLE Express (Gibco) for 5 minutes at 37 °C, followed by centrifugation at 500 × *g* for 5 minutes. The resulting cell clusters were combined with a viral suspension containing IntestiCult Organoid Growth Medium (Human) supplemented with 10 μM Y-27632 dihydrochloride (Millipore Sigma) and 8 μg/ml Polybrene (Millipore Sigma). The cells were transferred into a 24-well culture plate and centrifuged at 600 *× g* at 32 °C for 30 minutes. After 2 hrs of incubation at 37 °C, the cells were collected into 1.5 ml tubes and centrifuged at 300 *× g* for 5 minutes. Finally, the cells were embedded in 30 μl of Matrigel and cultured in 24-well plates with antibiotic-free IntestiCult Organoid Growth Medium (Human) supplemented with Y-27632 dihydrochloride. 3 days post-infection, the medium was replaced with culture medium containing 1 μg/ml puromycin. Organoid differentiation and downstream analyses, including immunofluorescence, apoptosis, and viability assays, were conducted as described above.

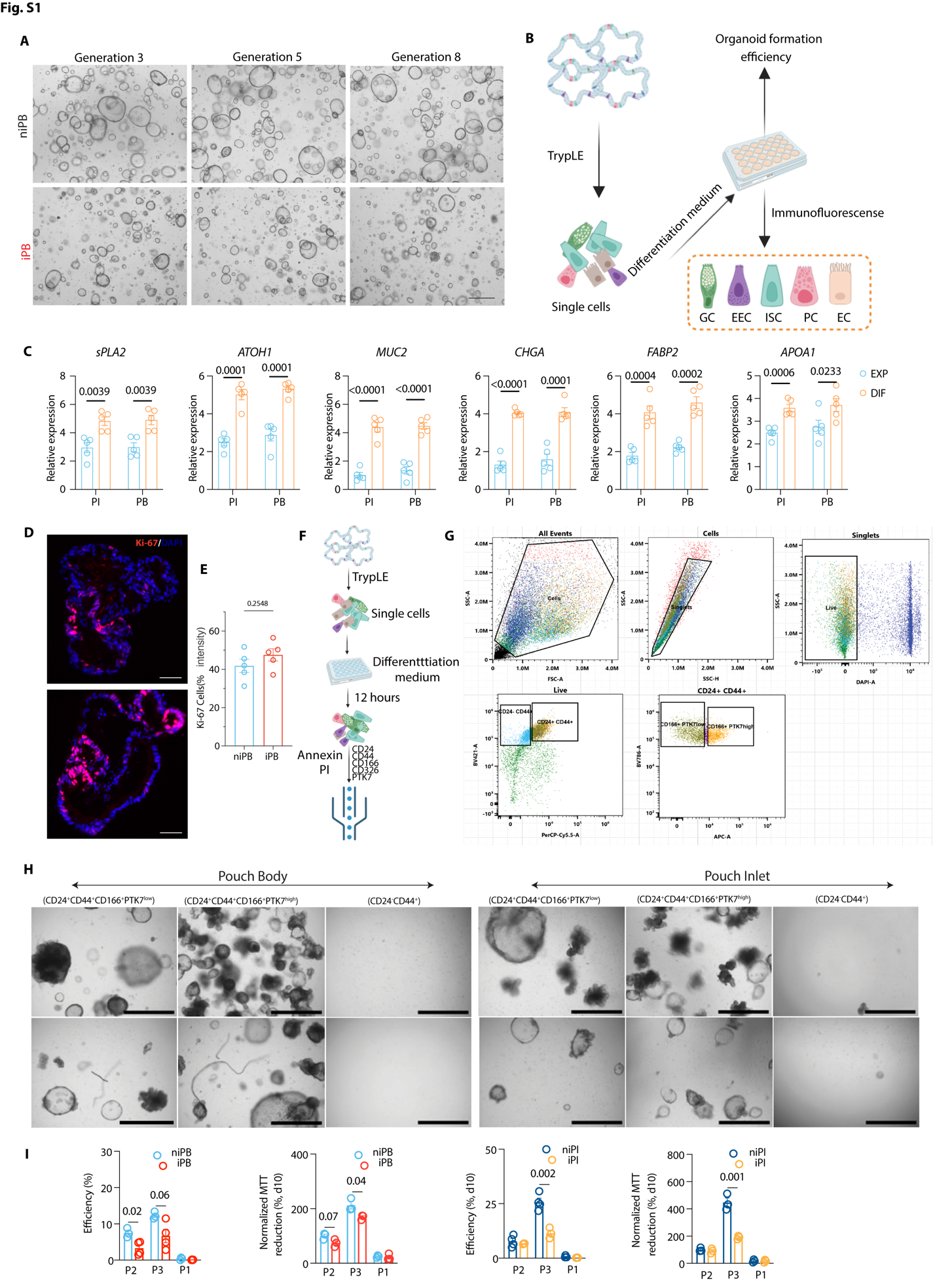

**Fig S1. Characterization of J-pouch derived organoid cultures.**

**(A)** Representative brightfield images of niPB and iPB at generations 3, 5, and 8. **(B)** Schematic representation of the organoid culture preparation for quantification of cell death and differentiation. GC: goblet cell; EEC: enteroendocrine cell; ISC: intestinal stem cell; PC: Paneth cell; EC: enterocyte cell. **(C)** RT-qPCR analysis of *sPLA2*, *ATOH1*, *MUC2*, *CHGA*, *FABP2*, and *APOA1* in pouch inlet (PI) and pouch body (PB) organoids cultured in expansion media (EXP) or differentiation media (DIF) for 7 days. **(D and E)** Representative immunofluorescence staining of Ki-67 (D) and quantification (E) in niPB and iPB organoids. Scale bars, 400 μm. **(F and G)** Schematic representation of the single stem cell preparation (F) and flow cytometric gating strategy (G) for ISC populations. **(H)** Representative images of organoids generated from indicated sorted cell populations in niPI, iPI, niPB, and iPB organoids. **(I)** Quantification of organoid generation efficiency and MTT assay from P1, P2, and P3 cells from niPI, iPI, niPB, and iPB organoids. For the MTT assay, P2 of niPB or niPI organoids was normalized to 100%. Dots represent individual samples; bars show mean ± SD. Statistical significance was assessed by one-way ANOVA with Tukey’s post hoc test. Exact P values are shown.

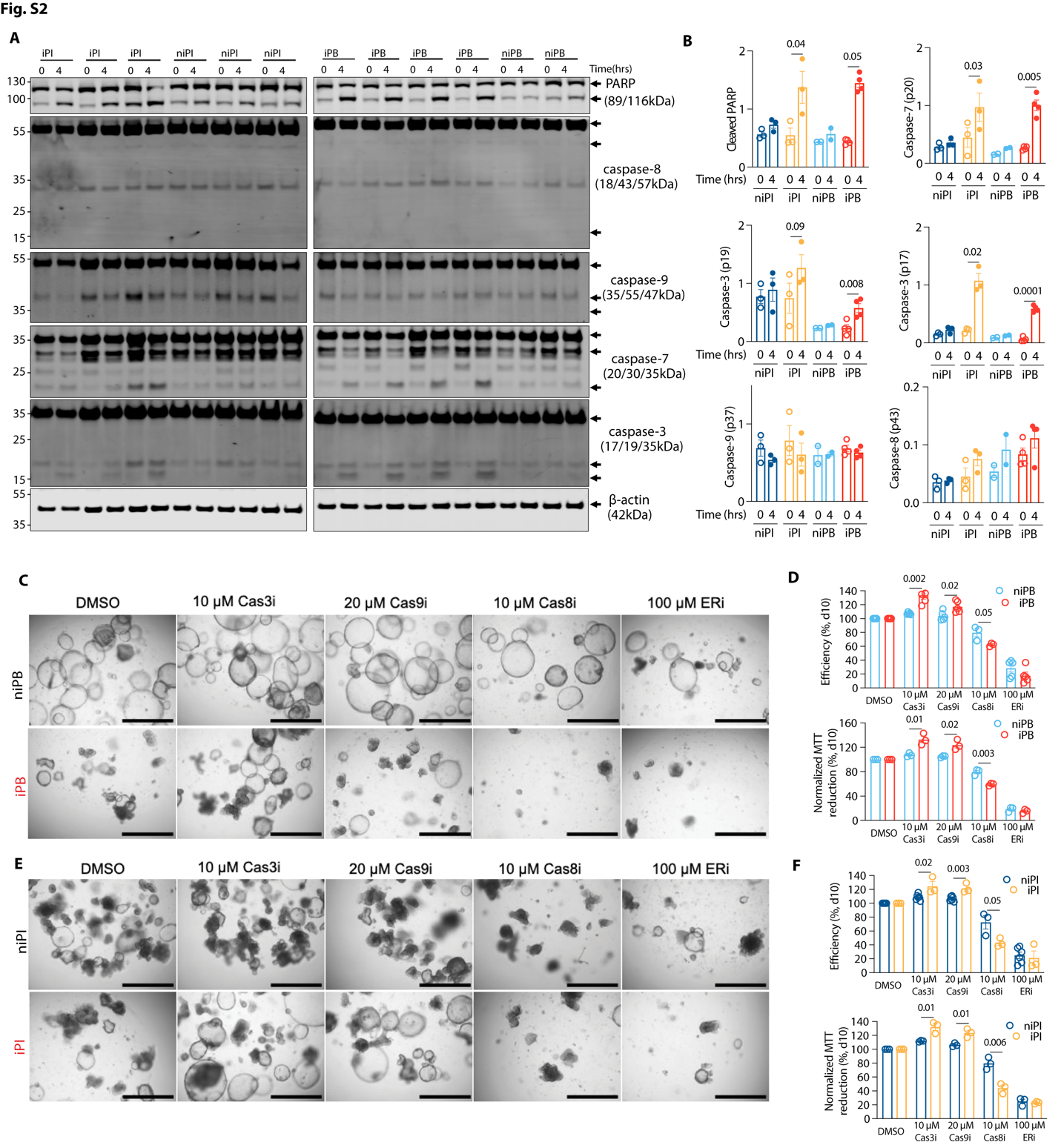

**Fig S2. Cell death of pouchitis organoids is associated with cleavage of caspase-3, 7, and 9.**

**(A)** Western blot analysis of PARP, caspase-8, caspase-9, caspase-3, caspase-7, and β-actin in cell lysates of representative niPI, iPI, niPB, and iPB organoids at 0 and 4 h. **(B)** Quantification of cleaved PARP, caspase-7 (p20), caspase-3 (p17 and p19), caspase-9 (p37) and caspase-8 (p43) band intensities relative to their respective pro-proteins in Western blot from (A). **(C-F)** Representative images (C and E) and quantification of organoid formation efficiency and MTT reduction (D and F) in niPB and iPB organoids treated with DMSO, 10 μM caspase-3 inhibitor (Cas3i, Z-DEVD-FMK), 20 μM caspase-9 inhibitor (Cas9i, Z-LEHD-FMK TFA), 10 μM caspase-8 inhibitor (Cas8i, Z-IETD-FMK), or 100 μM taurosodeoxycholic acid (TUDCA). Scale bars, 400 µm. Bars show the means ± SD. Statistical significance was assessed by two-way ANOVA with with Holm–Šidák correction for multiple comparisons. Exact P values are shown.

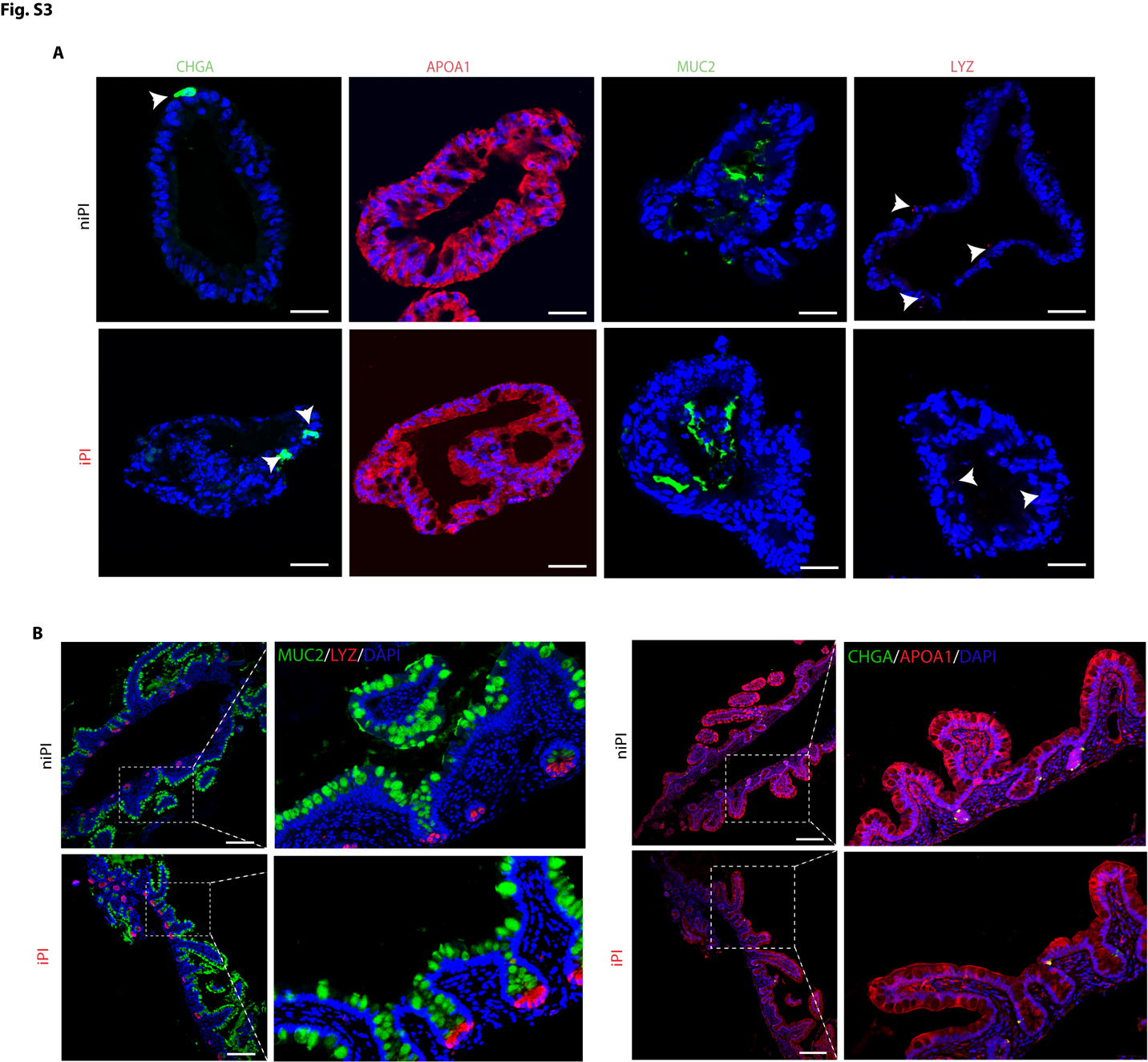

**Fig. S3. Pouchitis organoids display enhanced differentiation of secretory epithelial cells.**

**(A)** Immunofluorescence staining of organoids generated from inflamed and non-inflamed pouch inlet for MUC2, LYZ, CHGA, APOA1. Scale bars, 200 µm. **(B)** Immunofluorescence microscopy of J-pouch tissue sections from individuals with or without pouchitis stained for MUC2, LYZ, CHGA, and APOA1. Scale bars, 50 µm.

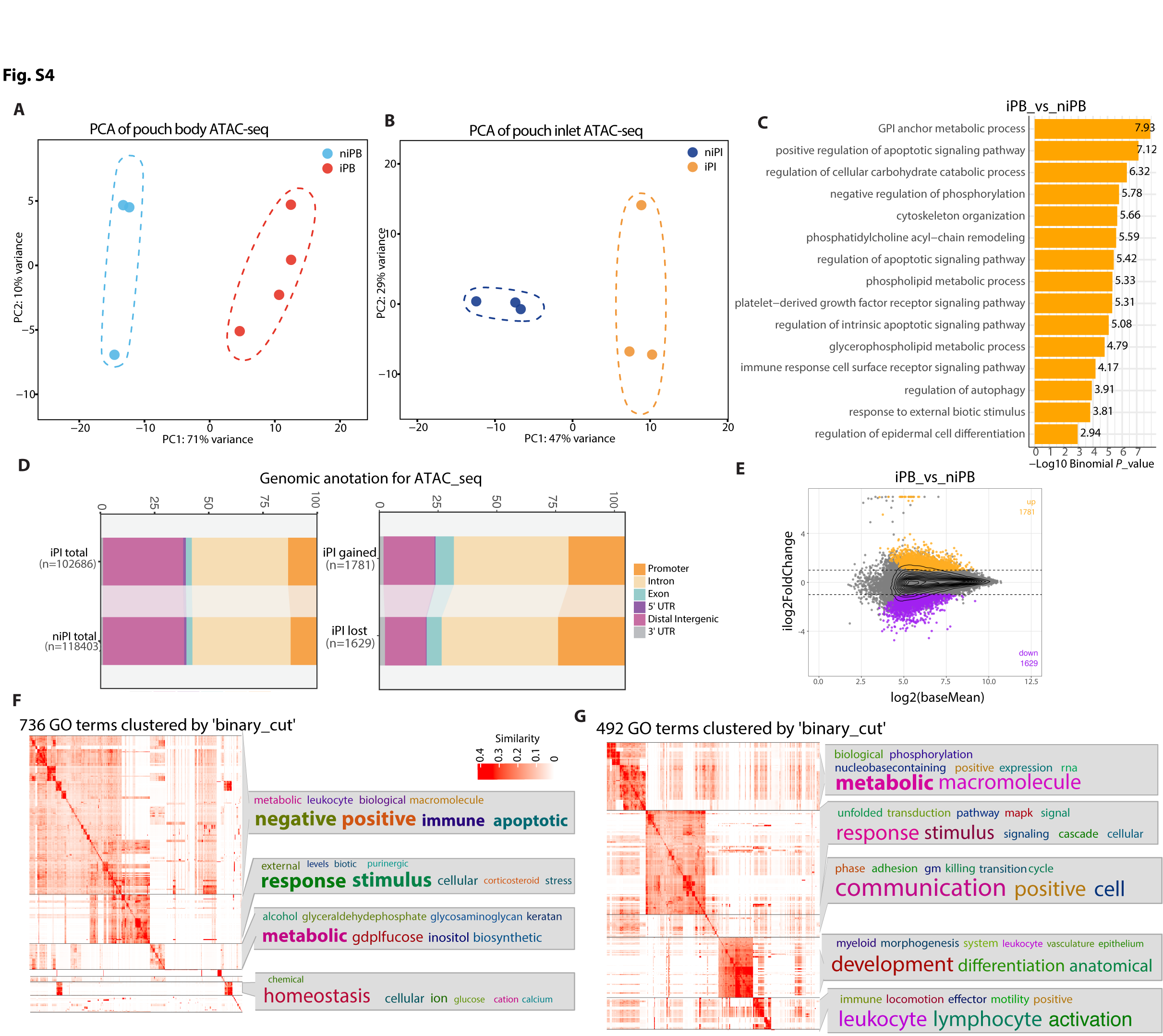

**Fig S4. ISCs from pouchitis organoids display alterations in chromatin accessibility.**

**(A and B)** Principal component analysis (PCA) of ATAC-seq profiles from ISCs isolated from pouch body (A) and pouch inlet (B) organoids. **(C)** Gene Ontology (GO) terms corresponding to biological processes associated with differentially regulated chromatin regions when comparing iPB versus niPB by Genomic Regions Enrichment of Annotations Tool (GREAT) analysis. **(D)** Genomic annotation of total accessible regions in iPI and niPI and regions with gained or lost accessibility in iPI versus niPI.**(E)** MA plots showing regions of lost and gained chromatin accessibility of ATAC-seq in pouch inlet. Significant increase or decrease was determined by DESeq2. Adjusted p-values were calculated using the Wald test with Benjamini-Hochberg correction (*P_adj_* < 0.05, fold change ≥ 1.5). **(F and G)** Gene Ontology (GO) biological process terms associated with differentially accessible chromatin regions identified by Genomic Regions Enrichment of Annotations Tool (GREAT) analysis in ISCs from niPI versus iPI gained (F) or lost (G). Similar GO terms were grouped based on semantic similarity and clustered using the binary-cut algorithm

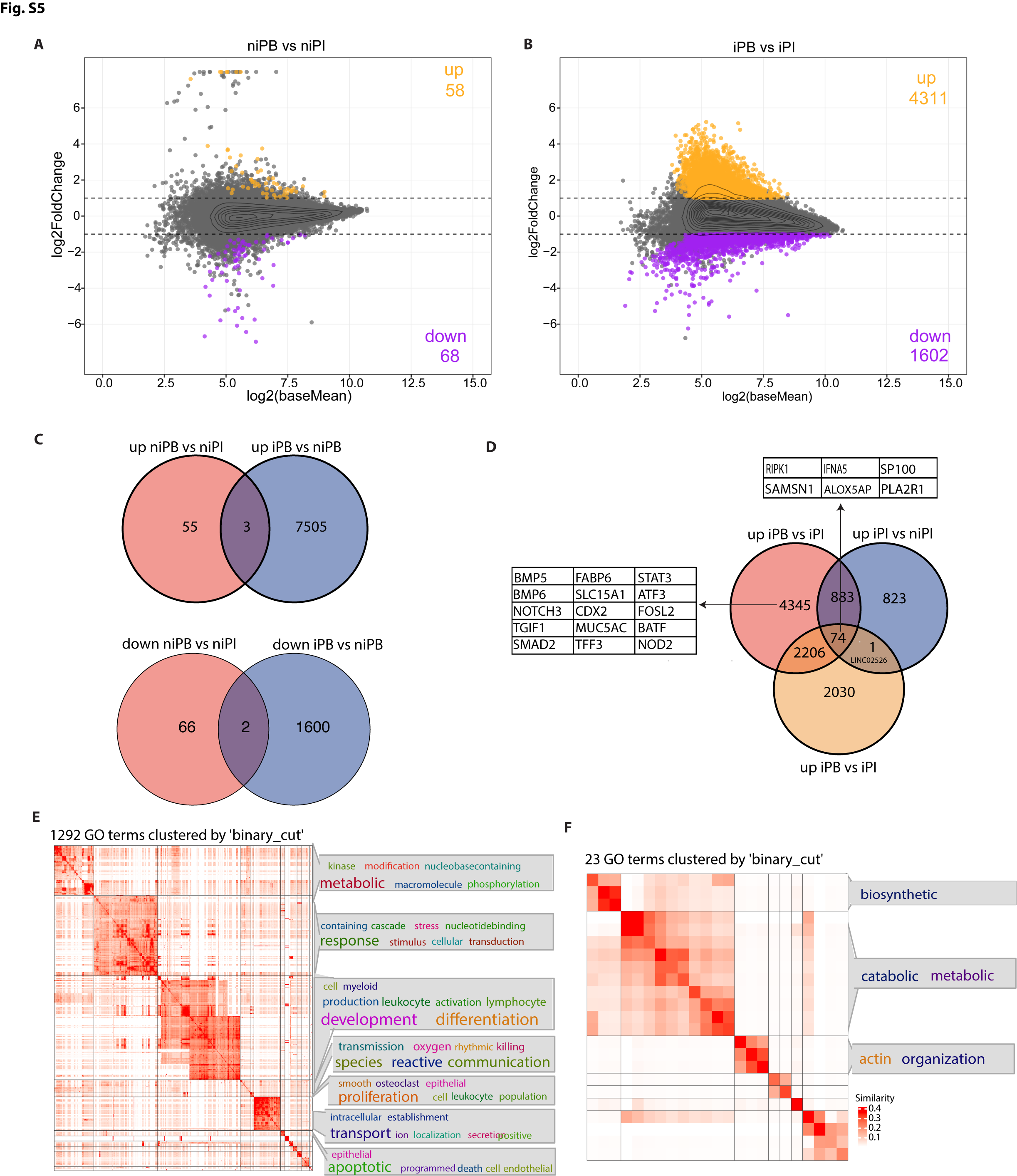

**Fig. S5. Inflammation-associated transcriptional remodeling is more pronounced in pouch body than pouch inlet intestinal stem cells.**

**(A and B)** MA plots showing differential accessibility identified by ATAC-seq in ISCs isolated from niPB versus niPI organoids (A) and iPB versus iPI (B). Regions with gained accessibility are shown in orange and regions with lost accessibility in purple (*P_adj_* < 0.05, |FC| > 1.5 **(C)** Venn diagrams comparing regions with gained accessibility (top) or lost accessibility (bottom) in niPB versus niPI and iPB versus niPB. The number of shared and unique regions in each comparison is indicated. **(D)** Venn diagram showing overlap among genes upregulated in iPB versus niPI, iPI versus niPI, and iPB versus iPI comparisons. Representative genes from selected overlapping and unique gene sets are shown. **(E)** Semantic clustering of Gene Ontology (GO) biological process terms associated with genes upregulated in niPB versus niPI. GO terms identified by enrichment analysis were grouped based on semantic similarity using the binary-cut algorithm. **(F)** Semantic clustering of GO biological process terms associated with genes upregulated in iPB versus iPI up-regulated (E) and down-regulated (F). Similar GO terms were grouped based on semantic similarity and clustered using the binary-cut algorithm

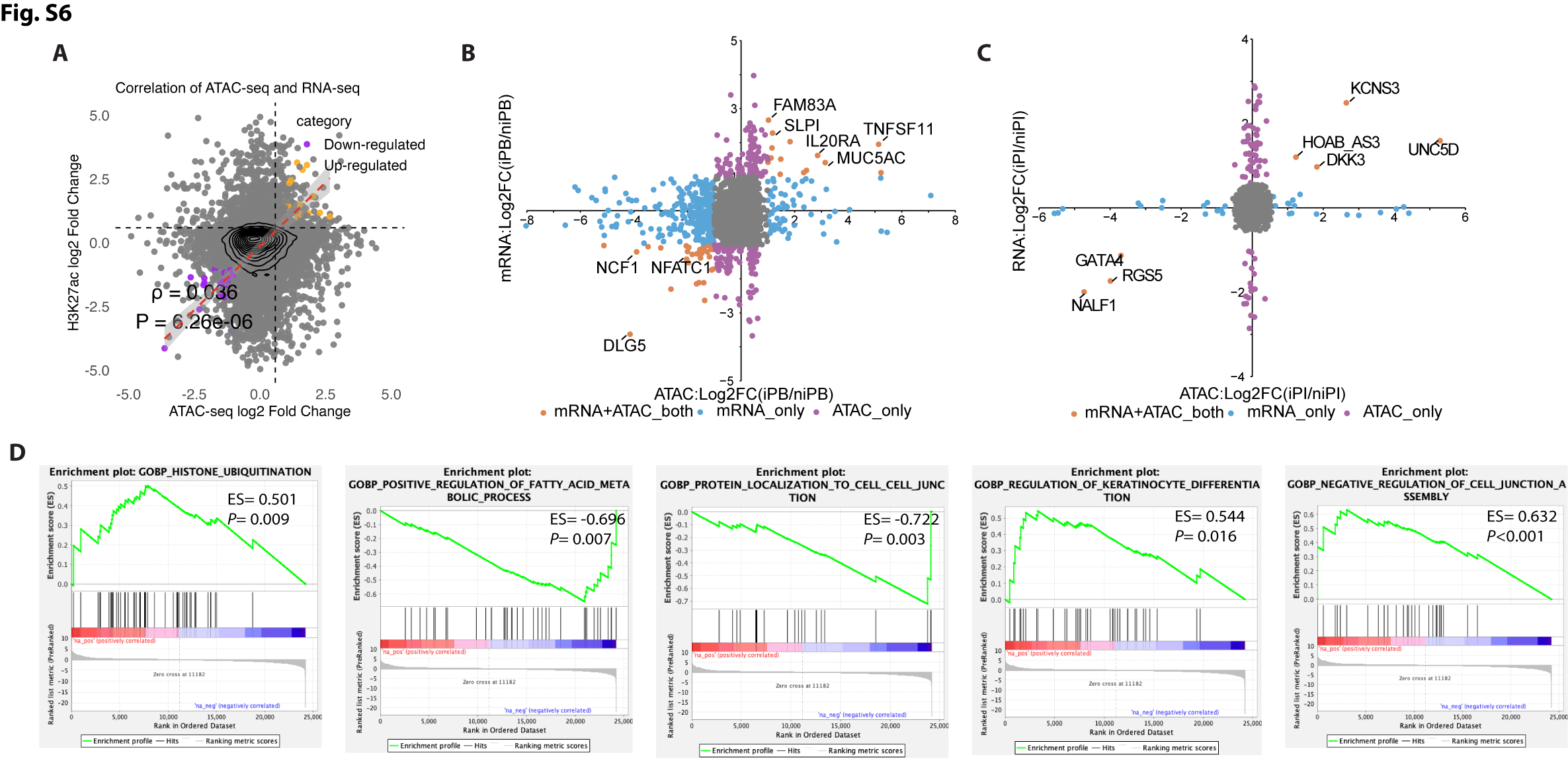

**Fig S6. Integrated ATAC-seq and RNA-seq analyses reveal correlations between chromatin remodeling and transcriptomes of iPB and niPB organoids.**

**(A)** Correlation between differential chromatin accessibility (ATAC-seq) and differential gene expression (RNA-seq) in pouch body ISCs. Each point represents a gene-associated chromatin region. Regions exhibiting coordinated changes in chromatin accessibility and transcription are highlighted. Spearman correlation coefficient (ρ) and corresponding P value are indicated. **(B and C)** Integration of ATAC-seq and RNA-seq datasets for pouch body (B) and pouch inlet (C) ISCs. Scatter plots display genes exhibiting concordant changes in chromatin accessibility and transcription (mRNA+ATAC), genes altered only at the chromatin level (ATAC-only), and genes altered only at the transcriptional level (mRNA-only). Representative genes are labeled. **(D)** Gene Set Enrichment Analysis (GSEA) plots showing significant enrichment of pathways, among genes with differential expression identified by RNA-seq.

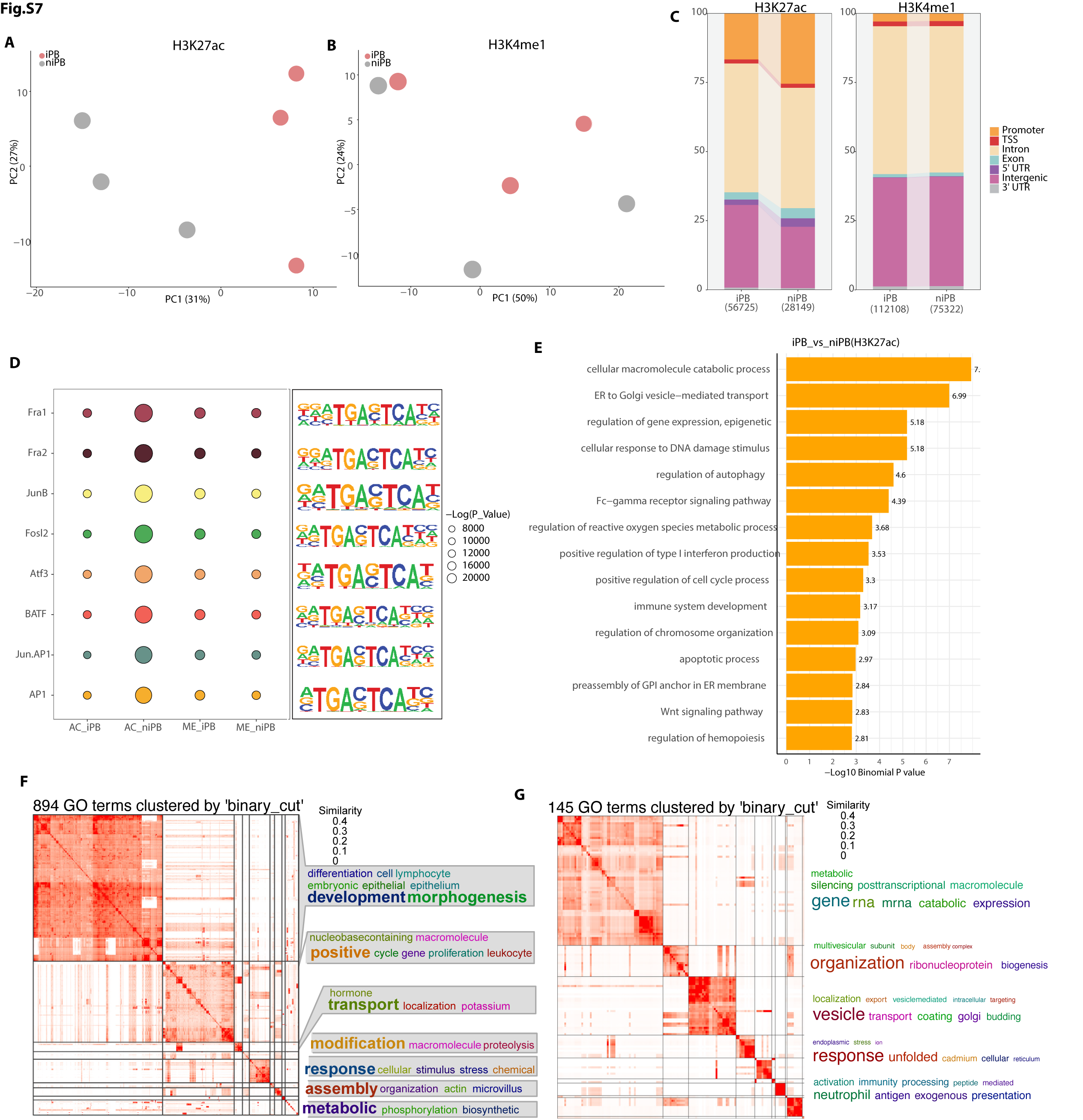

**Fig S7. ISCs from pouchitis organoids display alterations in histone modifications.**

**(A and B)** Principal Component Analysis (PCA) of H3K27ac (A) and H3K4me1 profiles in iPB and niPB ISCs. **(C)** Genomic annotation of total accessible regions of H3K27ac and H3K4me1 peaks across regions (promoter, TSS, exon, intron, UTR, distal intergenic) in iPB and niPB ISCs. **(D)** Dot plot and motif logos showing enriched transcription factor motifs on H3K27ac, including Jun/AP-1 and other key regulators. **(E)** GREAT analysis GO-terms of biological processes associated with gain and lost chromatin regions of H3K27ac. **(F and G)** Gene ontology (GO) terms corresponding to biological processes associated with chromatin regions that gained H3K27ac accessibility (F) or lost H3K27ac accessibility (G) in inflamed pouch body organoids compared with non-inflamed pouch body organoids. Similar GO terms were grouped based on semantic similarity and clustered using the binary-cut algorithm.

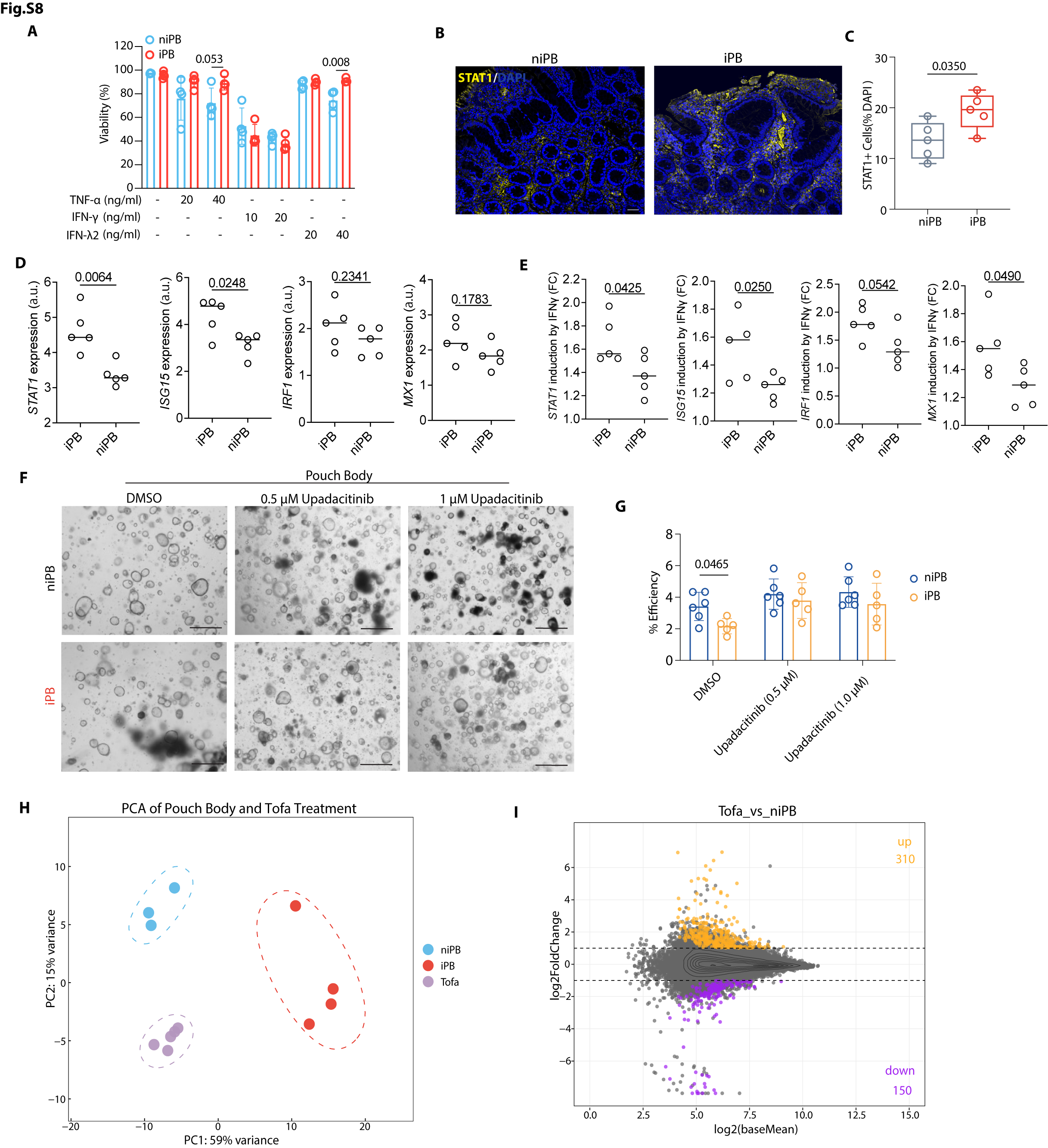

**Fig. S8. STAT1 activation and rescue of ISC dysfunction by JAK inhibition in the J-pouch.** **(A)** Organoid generation efficiency of organoids derived from niPB and inflamed pouch body (iPB) following treatment with TNF-α, IFN-γ, and IFN-λ2. **(B and C)** Representative immunofluorescence images (B) and quantification of STAT1 staining in niPB and iPB tissues (C). Scale bars, 50 μm. (D and E) RT-qPCR analysis of *STAT1, ISG15, IRF1*, and *MX1* expression in iPB and niPB organoids (D) and fold induction following IFN-γ stimulation (E). **(F and G)** Representative brightfield images (F) and quantification of organoid formation efficiency (G) in niPB and iPB organoids treated with upadacitinib (0.5 or 1 μM). **(H)** Principal component analysis (PCA) of ATAC-seq profiles from niPB, iPB, and tofacitinib-treated iPB (Tofa) ISCs. **(I)** MA plot showing differential chromatin accessibility in Tofa versus iPB ISCs. Statistical significance was assessed by two-way ANOVA with Holm–Šidák correction for multiple comparisons (A and G) and by an unpaired two-tailed Student’s t test (C). Exact P values are shown.

**Table S1. List of Antibodies, Chemicals, Peptides, Recombinant Proteins and Software Used in This Study**

| **REAGENT or RESOURCE** | **SOURCE** | **IDENTIFIER** |
| --- | --- | --- |
| Antibodies | | |
| Mouse anti-Chr-A | Santa Cruz Biotechnology | Cat#: sc-393941 |
| Mouse anti-Mucin 2 | Santa Cruz Biotechnology | Cat#: sc-515032 |
| ApoA1 Polyclonal Antibody | Invitrogen | Cat#: PA5-88109 |
| Anti-lysozyme antibody | abcam | Cat#: ab108508 |
| anti-human CD44 Antibody, BV421 | BioLegend | Cat#: 103039 |
| anti-human PTK7(CCK-4), APC | Miltenyi Biotec | Cat#: 130-099-660 |
| anti-human CD24, PerCP-Cy5.5 | BioLegend | Cat#: 311116 |
| Anti-Human CD326, BV786 | BD Biosciences | Cat#: 565685 |
| CD166 Monoclonal Antibody | Invitrogen | Cat#: 12-1668-42 |
| Annexin V, FITC | BD Biosciences | Cat#: 556419 |
| H3K27Ac Antibody | Cell Signaling Technology | Cat#: 8173S |
| H3K4Me1 Antibody | Cell Signaling Technology | Cat#: 5326S |
| Rabbit monoclonal anti c-Jun | Cell Signaling Technology | Cat#: 9165S |
| Rabbit monoclonal anti p-c-Jun | Cell Signaling Technology | Cat#:3270S |
| Rabbit monoclonal anti c-Fos | Cell Signaling Technology | Cat#: 74620S |
| Rabbit monoclonal anti GAPDH | Cell Signaling Technology | Cat#: 2118S |
| Rabbit monoclonal anti FRA1 | Cell Signaling Technology | Cat#: 5281S |
| Rabbit monoclonal anti BATF | Cell Signaling Technology | Cat#: 8638S |
| Rabbit monoclonal anti OLMF4 | Cell Signaling Technology | Cat#: 14369S |
| Mouse monoclonal anti IKKα | Cell Signaling Technology | Cat#:11930T |
| Rabbit monoclonal anti IKKβ | Cell Signaling Technology | Cat#:8943T |
| Rabbit monoclonal anti p65 | Cell Signaling Technology | Cat#: 3033S |
| Rabbit monoclonal anti IKKα/β | Cell Signaling Technology | Cat#: 2697S |
| Rabbit monoclonal anti SAPK/JNK | Cell Signaling Technology | Cat#: 99252S |
| Rabbit monoclonal anti β-Catenin | Cell Signaling Technology | Cat#: 8441SF |
| Chemicals, peptides, and recombinant proteins | | |
| ProLong Glass Antifade Mountant Stain | ThermoFisher | Cat#: P36934 |
| 16% paraformaldehyde | Sigma-Aldrich | Cat#: 15710S |
| Triton X-100 | Sigma-Aldrich | Cat#: T8787 |
| Tween-20 | Sigma-Aldrich | Cat#: P9416 |
| DMEM/F12 | Gibco | Cat#: 11320033 |
| TrypLE express | Gibco | Cat#: 12604013 |
| Cell Recovery Medium | Corning | Cat#: 354253 |
| TrypLE express | Gibco | Cat#: 12605010 |
| Y-27632 | Sigma-Aldrich | Cat#: Y05030 |
| Human Wnt-3a Protein | R&D Systems | Cat#: 5036-WN-010 |
| Human Noggin Protein | R&D Systems | Cat#:6057-NG-100 |
| Human R-Spondin 1 Protein | R&D Systems | Cat#:4645-RS-100 |
| Human EGF | R&D Systems | Cat#: 236-EG |
| Human FGF | Peprotech | Cat#: 100-18B |
| Human IGF-1 | BioLegend | Cat#: 590908 |
| Gastrin-Leu15 | Sigma-Aldrich | Cat#: G9145 |
| N-2 Supplement | Gibco | Cat#: 17502048 |
| A 83-01 | BioGems | Cat#: 9094360 |
| N-Acetyl-L-cysteine | Sigma-Aldrich | Cat#: A9165 |
| B27 | Gibco | Cat#: |
| GlutaMAX™ Supplement | Gibco | Cat#: 35050061 |
| Gentamicin | Gibco | Cat#: 15750060 |
| Gentle Cell Dissociation Reagent | Stemcell | Cat#: 100-0485 |
| IntestiCultÔ Organoid GrowthMedium (Human) | Stemcell | Cat#: 06010 |
| Penicillin-Streptomycin | Sigma-Aldrich | Cat#: P4333 |
| Gentamicin (50 mg/mL) | Gibco | Cat#: 15750060 |
| Bovine Serum Albumin | Sigma-Aldrich | Cat#: A7030-10G |
| Advanced DMEM/F-12 | Gibco | Cat#: 12634028 |
| NP-40 lysis buffer | Thermo Fisher | Cat#: J60766.AK |
| Sucrose | Sigma-Aldrich | Cat#: S0389 |
| Puromycin | Sigma-Aldrich | Cat#: P8833 |
| BlockAid™ Blocking Solution | Thermo Fisher | Cat#: B10710 |
| AMPure XP Reagent | BECKMAN BCOUKTER | Cat#: A63880 |
| Intercept® (TBS) Blocking Buffer | Licor Bio | Cat#: 927-60001 |
| Lenti-X™ Concentrator | TaKaRa | Cat#: 631231 |
| shRNA control in pGFP-C-shLenti shRNA Vector | OriGene | Cat#: TR3002 |
| NEG-50Ô Frozen Section Medium | Epredia | Cat#: 6502 |
| Critical commercial assays | | |
| CUT&RUN Assay Kit | Cell Signaling Technology | Cat#: 86652S |
| DNA Library Prep Kit | Cell Signaling Technology | Cat#: 56795S |
| Multiplex Oligos for Illumina Systems | Cell Signaling Technology | Cat#: 47538S |
| Lentiviral Packaging Kits | OriGene | Cat#: TR30037 |
| c-Jun (JUN) Human shRNA Plasmid Kit | OriGene | Cat#: TL320297 |
| QIAGEN Plasmid Kits | Qiagen | Cat#: 12143 |
| BCA protein assay kit for low concentrations | abcam | Cat#: ab207002 |
| Software and algorithms | | |
| R-Studio | Posit software | <https://posit.co/download/rstudio-desktop/> |
| R | The R Foundation | <https://www.r-project.org/> |
| MACS2 | Zhang et al., 2008; Feng et al., 2012 | <https://github.com/macs3-project/MACS> |
| Deeptools | Ramírez et al., 2016 | <https://deeptools.readthedocs.io/en/latest/> |
| limma | [Ritchie et al., 2015](https://www.sciencedirect.com/science/article/pii/S1934590921002861?casa_token=glBpkotn1AcAAAAA:miRihmYz_zi5xJ0bQ_ku4Vl-taTdTvwkmwEe8WSZZDmvb8LGDFuWM8ZpNQeMB5fqW9U7AhE#bib60) | [https://kasperdanielhansen.github.io/genbioconductor/html/](https://kasperdanielhansen.github.io/genbioconductor/html/limma.html)limma.html |
| DESeq2 | Love et al., 2014 | <https://www.bioconductor.org/packages/devel/bioc/vignettes/DESeq2/inst/doc/DESeq2.html> |
| GREAT | McLean et al., 2010 | <https://great.stanford.edu/great/public/html/> |
| Profileplyr | Carroll and Barrows, 2020 | <https://www.bioconductor.org/packages/devel/bioc/vignettes/profileplyr/inst/doc/profileplyr.html> |
| HOMER | Heinz et al., 2010 | <http://homer.ucsd.edu/homer/index.html> |
| MotifMatchr | Schep, 2019 | <https://github.com/GreenleafLab/motifmatchr> |
| TFBSTools | Tan and Lenhard, 2016 | <https://bioconductor.org/packages/release/bioc/vignettes/TFBSTools/inst/doc/TFBSTools.html> |
| TOBIAS | Bentsen et al., 2020 | <https://github.com/loosolab/TOBIAS> |
| Bowtie2 | Langmead et al., 2009 | <https://bowtie-bio.sourceforge.net/bowtie2/index.shtml> |
| Samtools | Li et al., 2009 | <https://www.htslib.org/> |
| SEACR | [Meers et al., 2019](https://www.sciencedirect.com/science/article/pii/S1934590921002861?casa_token=glBpkotn1AcAAAAA:miRihmYz_zi5xJ0bQ_ku4Vl-taTdTvwkmwEe8WSZZDmvb8LGDFuWM8ZpNQeMB5fqW9U7AhE#bib40) | [https://seacr.fredhutch.org](https://seacr.fredhutch.org/) |
| MotifDb | Shannon and Richards, 2019 | <https://github.com/tmuetze/MotifDb> |
| GSEA | Subramanian et al., 2005; Mootha et al., 2003 | <https://www.gsea-msigdb.org/gsea/index.jsp> |
| Genrich | John M. Gaspar | <https://github.com/jsh58/Genrich> |
| Prism 10 | GraphPad Software | <https://www.graphpad.com/features> |
| Fiji | NIH | <https://imagej.net/software/fiji/downloads> |
| FlowJo | BD and FlowJo | <https://www.flowjo.com/> |

**Table S2.** Information for patients with Crohn’s disease, ulcerative colitis, and ulcerative colitis with IPAA. M, male; F, female.

| **Name** | **Sex** | **Disease** | **Location** | **Endoscopic Disease Activity** | **Processed Sample Type** |
| --- | --- | --- | --- | --- | --- |
| niPI1 | F | Inflammatory bowel disease with IPAA | Pouch inlet | Non-Inflamed | Organoids and tissue section |
| niPI2 | F |  |  |  | Organoids and tissue section |
| niPI3 | F |  |  |  | Organoids and tissue section |
| niPI4 | M |  |  |  | Organoids and tissue section |
| niPI5 | F |  |  |  | Organoids and tissue section |
| niPI6 | F |  |  |  | Organoids and tissue section |
| niPI7 | M |  |  |  | Organoids and tissue section |
| niPI8 | M |  |  |  | Organoids and tissue section |
| niPI9 | M |  |  |  | Organoids and tissue section |
| niPI10 | M |  |  |  | Organoids and tissue section |
| niPI11 | F |  |  |  | Organoids and tissue section |
| IPI1 | M |  |  | Inflamed | Organoids and tissue section |
| IPI2 | F |  |  |  | Organoids and tissue section |
| IPI3 | F |  |  |  | Organoids and tissue section |
| IPI4 | M |  |  |  | Organoids and tissue section |
| IPI5 | F |  |  |  | Organoids and tissue section |
| iPB1 | F |  | Pouch body | Non-inflamed | Organoids and tissue section |
| iPB2 | F |  |  |  | Organoids and tissue section |
| iPB3 | F |  |  |  | Organoids and tissue section |
| iPB4 | F |  |  |  | Organoids and tissue section |
| iPB5 | M |  |  |  | Organoids and tissue section |
| iPB6 | M |  |  |  | Organoids and tissue section |
| iPB7 | M |  |  |  | Organoids and tissue section |
| iPB8 | M |  |  |  | Organoids and tissue section |
| iPB9 | F |  |  |  | Organoids and tissue section |
| iPB1 | M |  |  | Inflamed | Organoids and tissue section |
| iPB2 | F |  |  |  | Organoids and tissue section |
| iPB3 | M |  |  |  | Organoids and tissue section |
| iPB4 | F |  |  |  | Organoids and tissue section |
| iPB5 | M |  |  |  | Organoids and tissue section |
| iPB6 | M |  |  |  | Organoids and tissue section |
| iPB7 | M |  |  |  | Organoids and tissue section |

**Table S3. Base line characteristics of the patients.**

|  | **Pouch (n=22)** |
| --- | --- |
| Age (mean; range) | 40.71 (23-73) |
| Sex—male | 12 (57.14%) |
| Disease duration (mean; range) | 16.10 (1-37) |
| Proctitis (E1) | NA |
| Left-sided (E2) | NA |
| Pancolitis (E3) | NA |
| A1 (dx<16y) | NA |
| A2 (dx 17-40y) | NA |
| A3 (dx>40y) | NA |
| L1 (ileal) | NA |
| L2 (colonic) | NA |
| L3 (ileocolonic) | NA |
| B1 (non-stricturing, non-penetrating) | NA |
| B2 (stricturing) | NA |
| B3 (penetrating) | NA |
| P (perianal disease) | NA |
| CRP**** (mg/L) | 3.4 (0-14.6) |
| Fecal calprotectin (µg/g) | NA |
| Endoscopic Pouch Disease Activity Index (ePDAI) (mean; range) | 1.19 (0-4) |
| Endoscopic Mayo Score (UC) (mean; range) | NA |
| Simple Endoscopic Score for Crohn's Disease (SES-CD) (mean; range) | NA |

***NA, not applicable; ****CRP, C-reactive protein. **Table S4. Base line characteristics of the patients.**

**Table S4.** Primer sequences used for RT-qPCR. All primer sequences are shown in the 5′ to 3′ direction.

| Gene | Forward primer (5′→3′) | Reverse primer (5′→3′) |
| --- | --- | --- |
| *MUC2* | CTGGACTGGTTGGACGATC | TGCACCTCTTGTAATACTCAGC |
| *JUN* | AGCCCAAACTAACCTCACG | TGCTCTGTTTCAGGATCTTGG |
| *HES1* | AACGCAGTGTCACCTTCC | TCAAGTTCCTGTTTAGAGTCCG |
| *CHGA* | ATGTTTTGAGACACTCCGAGG | GAGTTCATCTTCAAAACCGCTG |
| *NEUROD1* | CCAGGGTTATGAGACTATCACTG | TCCTGAGAACTGAGACACTCG |
| *NEUROG3* | GGCAGTCTGGCTTTCTCAG | GGAGAAGCAGAAGGAACAAGTG |
| *STAT1* | CACCTACGAACATGACCCTATC | GCTGTCTTTCCACCACAAAC |
| *IRF1* | GTGTGGATCTTGCCACATTTC | CCGAGCAAGGCACTGTATAA |
| *APOA1* | TGTGTCCCAGTTTGAAGGC | CTCCTTTTCCAGGTTATCCCAG |
| *ATOH1* | AGGAAAACAGCAAAACTTCGC | AGTCACTGTAATGGGAATGGG |
| *LYZ* | TGATCCACAAGGCATTAGAGC | GAACATACTGACGGACATCTCTG |
| *GAPDH* | ACATCGCTCAGACACCATG | TGTAGTTGAGGTCAATGAAGGG |
| *FOS* | TTGTGAAGACCATGACAGGAG | CCATCTTATTCCTTTCCTTTCGG |
| *MX1* | GAAGATAAGTGGAGAGGCAAGG | CTCCAGGGTGATTAGCTCATG |
| *PLA2G2A (sPLA2)* | CATTCACCTGCCCTGTCTC | CACTCCTGCTCCCCTTAAATAG |
| *FABP2* | GACTCAACTGAAATCATGGCG | TCAAATTGTCATGAGCTGCAAG |
